# Pyruvate kinase activates SARM1 to exacerbate axonal degeneration in diabetic peripheral neuropathy

**DOI:** 10.1101/2025.05.28.656718

**Authors:** Jingxuan Liu, Wenxi Li, Chenyu Li, Yuefeng jiang, Wenjie Zhu, Zhe Zhang, Yongjuan Zhao, Jinzhong Lin, Wei Yu

## Abstract

Diabetic peripheral neuropathy (DPN) is a prevalent and disabling complication of diabetes, characterized by progressive axonal degeneration. However, the molecular link between hyperglycemia and axon injury remains unclear. Here, we identify pyruvate kinase M (PKM) as a direct metabolic activator of the NADase SARM1 under high-glucose conditions. Proteomic and biochemical analyses reveal that PKM binds the TIR domain of SARM1 via its C-terminal region, allosterically enhancing NADase activity independently of PKM’s glycolytic role. In dorsal root ganglion (DRG) neurons, hyperglycemia strengthens the PKM–SARM1 interaction, driving NAD⁺ depletion, axonal fragmentation, and sensory dysfunction. Genetic depletion of PKM protects against streptozotocin-induced neuropathy, preserving nerve fiber density, restoring NAD⁺ levels, and alleviating mechanical allodynia. Most notably, we developed Pep-SP1, a competitive inhibitory peptide derived from residues 645–655 of the SARM1 TIR domain, which selectively disrupts the PKM–SARM1 interaction without impairing PKM metabolism or SARM1 catalytic activity. Systemic delivery of Pep-SP1 attenuates axonal degeneration and improves sensory outcomes in diabetic mice. By targeting a disease-potentiating interface, we expand therapeutic strategies beyond catalytic and allosteric inhibition, offering a mechanistically distinct avenue for axon protection with broad relevance to metabolic and neurodegenerative disorders.

## Introduction

Diabetic peripheral neuropathy (DPN) affects over half of individuals with both type 1 and type 2 diabetes and represents a major cause of morbidity^1^, manifesting as sensory loss, neuropathic pain and foot ulceration that can progress to limb amputation and mortality^1^. At the earliest stages, DPN is defined by distal axonopathy of peripheral sensory fibers^2^, yet the molecular cues that translate chronic hyperglycaemia into axonal degeneration remain poorly defined.

Sterile alpha and Toll/interleukin-1 receptor motif-containing protein 1 (SARM1) has emerged as the executioner of programmed axon degeneration^34567^: its intrinsic NAD^+^ hydrolase activity precipitates rapid NAD^+^ depletion and structural disassembly of injured axons^378^. Structural and biochemical studies indicate that SARM1 is allosterically^8^ inhibited by physiological NAD^+^ concentration^79^ and that diverse insults-axotomy, oxidative stress or metabolic challenge-trigger its activation^5^. However, how diabetic hyperglycaemia engages SARM1 to drive the neurodegenerative cascade in DPN has not been elucidated^4^.

Pyruvate kinase M (PKM) catalyses the final, ATP-generating step of glycolysis^1011^ and exists as two splice isoforms, PKM1 and PKM2, with distinct tissue distributions and regulatory properties^121314^. Beyond its metabolic role, PKM has been implicated in diabetic complications such as insulin resistance and microvascular dysfunction^131516^, but its contribution to axonal integrity under hyperglycaemic stress is unknown.

Here, we report that PKM directly binds the TIR domain of SARM1 via its C-terminal region and, independently of its kinase activity, allosterically potentiates SARM1’s NADase function under high-glucose conditions. In dorsal root ganglion neurons exposed to hyperglycaemia, enhanced PKM-SARM1 interaction precipitates NAD^+^ loss and axon fragmentation. Genetic ablation of PKM in mice preserves intraepidermal nerve fibers, restores DRG NAD^+^ levels and ameliorates mechanical allodynia in streptozotocin-induced DPN. Moreover, a peptide designed to occlude the PKM-SARM1 interface disrupts this interaction in vivo and rescues neuropathic phenotypes. These findings define a previously unrecognized, metabolism-driven mechanism of SARM1 activation and nominate the PKM-SARM1 axis as a tractable target for preventing axonal degeneration in diabetic neuropathy.

## Results

### PKM Directly Binds SARM1 and Allosterically Enhances Its NADase Activity

Given SARM1’s central role in axonal degeneration, we sought to uncover its upstream metabolic regulators by performing affinity purification from Expi293F cells coupled with mass spectrometry. Among the top interactors, the glycolytic enzyme pyruvate kinase M (PKM) and the mitochondrial channel VDAC1 were identified (Fig. 1A). Given PKM’s central role in neuronal energy metabolism^17^, we focused subsequent validation on this interaction.

**Figure 1.**
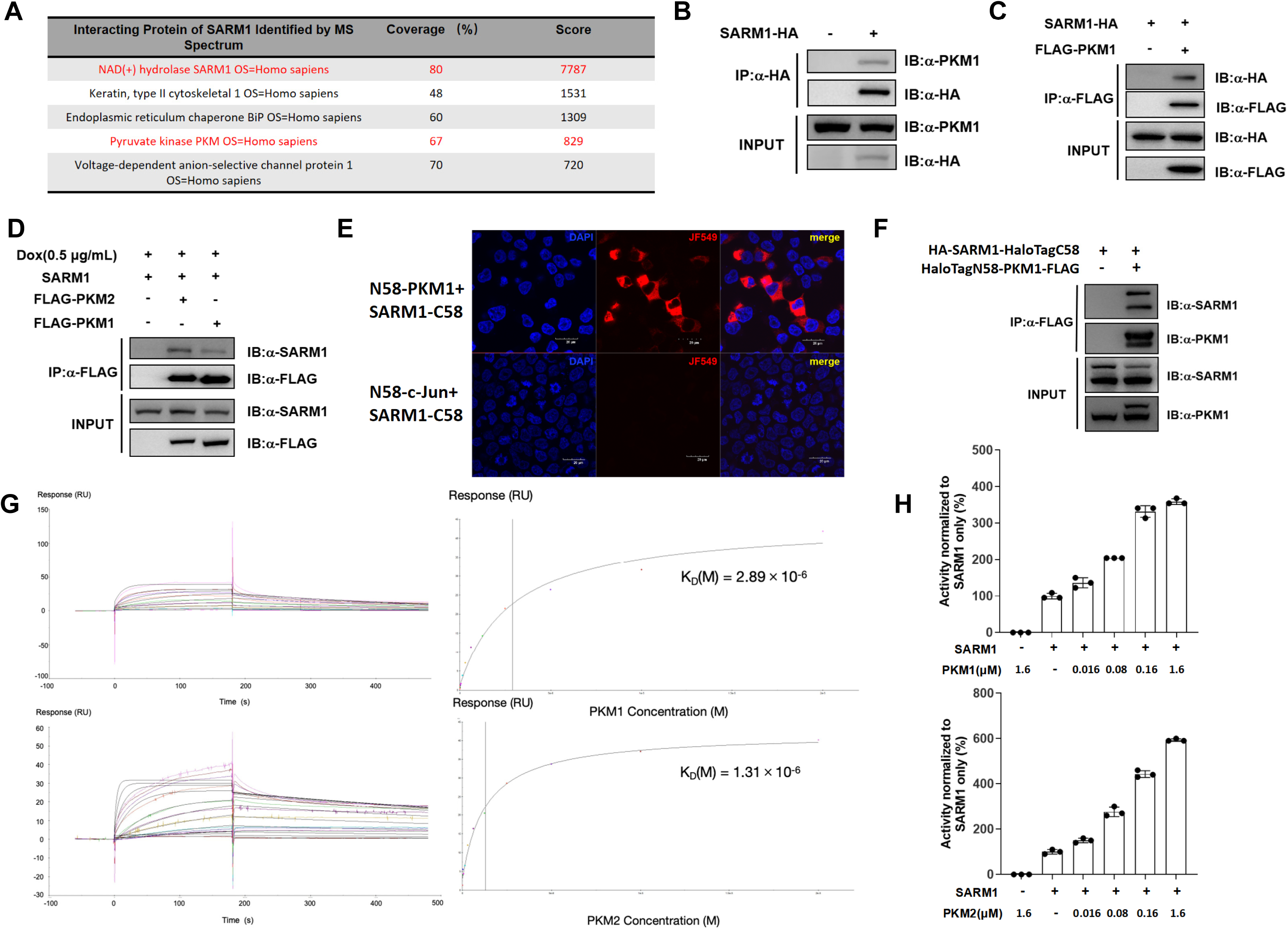
PKM directly binds SARM1 and allosterically enhances its NADase activity. (**A**) SARM1-interacting proteins were identified by mass spectrometry. The identified proteins are shown. PKM coverage was 67%, indicating that 67% of its amino acid sequence was detected by mass spectrometry. The score for PKM was 829, which reflects the total peptide match score and indicates a high confidence in the identification of PKM. (**B**) Endogenous PKM1 interacts with overexpressed SARM1-HA in HT22 cells. Lysates from HT22 cells transfected with SARM1-HA plasmid (2 μg, 24 h) were immunoprecipitated with anti-HA antibody. Western blotting with anti-PKM1 detected endogenous PKM1 (58 kDa) in the immunoprecipitate. Input lanes (10% total lysate) showed endogenous PKM1 and SARM1-HA expression levels. **(C**) Overexpressed SARM1-HA specifically binds FLAG-PKM1 in HT22 cells. SARM1-HA (2 μg) and FLAG-PKM1 (2 μg) plasmids were co-expressed for 24 h. Anti-FLAG immunoprecipitate probed with anti-HA confirms SARM1-HA/FLAG-PKM1 interaction. SARM1-HA: 72 kDa; FLAG-PKM1: 58 kDa. Input controls showed transfection efficiency. (**D**) Dox-inducible tag-free SARM1 interacts with PKM1 and PKM2 in HT22 cells. Dox (0.5 μg/mL, 16 h)-induced expression of tag-free SARM1 in 293T cells. Lysates were immunoprecipitated with anti-FLAG antibody, followed by western blotting with anti-SARM1. PKM1 (58 kDa) and PKM2 (60 kDa) bands indicate isoform-specific interactions. Input lanes (10% total lysate) demonstrated expression levels. (**E**) Bimolecular Fluorescence Complementation (BiFC) validates SARM1-PKM1 interaction. HaloTag-N58-PKM1 and SARM1-HaloTag-C58 were co-transfected into 293T cells for 24 h. Fluorescent signals (red, JF549) indicated reconstituted fluorescence due to SARM1-PKM1 interaction. Nuclei were counterstained with DAPI (blue). Negative controls: co-transfection of HaloTag-N58-c-Jun and SARM1-HaloTag-C58 showed no fluorescence (lower panel). Images were acquired using a confocal microscope (40× water objective). Scale bar: 20 μm. (**F**) Co-IP confirms the interaction between BiFC fusion proteins. Lysates from 293T cells co-expressing HaloTag-N58-PKM1-FLAG and SARM1-HaloTag-C58 for 24 h were immunoprecipitated with anti-FLAG antibody. Western blotting with anti-SARM1 detected SARM1-HaloTag-C58 in the immunoprecipitate. Input lanes (10% total lysate) confirmed protein expression levels. (**G**) The binding affinity between SARM1 and PKM1/2 was measured by surface plasmon resonance (left), affinity of SARM1 to PKM1 was 2.89 μM, and to PKM2 was 1.31 μM (right). (**H**) PKM1/PKM2 enhances SARM1 NADase activity in vitro. Recombinant SARM1 (6 µg/mL) was incubated with NAD^+^ (2000 μM), NMN (500 µM) and PC6 (50 µM) in the presence or absence of PKM1/PKM2 (0.016, 0.08, 0.16, 1.6 µM). PKM isoforms significantly accelerated NAD^+^ degradation. Data normalized to control (100% activity = SARM1 without PKM1 or PKM2). Bars represent standard error.

Endogenous PKM1 co-immunoprecipitated with HA-tagged SARM1 in HT22 neurons, confirming physiological binding (Fig. 1B). Robust reciprocal co-IPs in both HT22 and 293T cells co-expressing FLAG-PKM1 and SARM1-HA further validated this association (Fig. 1C, Fig. S1A-B). PKM2 bound SARM1 with comparable affinity (Fig. S1C-D), and doxycycline-inducible, untagged SARM1 retained interaction with FLAG-PKM1/2, ruling out epitope-tag artifacts (Fig. 1D).

To visualize binding in live cells, we adapted a split Halo-Tag complementation assay. Co-expression of HaloTag-N58-PKM1 and SARM1-HaloTag-C58 in 293T cells yielded punctate JF549 fluorescence, whereas the HaloTag-N58-c-Jun control showed no signal (Fig. 1E). Parallel co-IPs confirmed specific pulldown of both fusion partners alongside their endogenous counterparts (Fig. 1F). Surface plasmon resonance demonstrated direct PKM1-SARM1 and PKM2-SARM1 interactions with K_D_ values of 2.89 µM and 1.31 µM, respectively (Fig. 1G).

Given SARM1’s NADase function as the effector of axon degeneration, we tested whether PKM modulates this enzymatic activity. In vitro NAD^+^-hydrolysis assays revealed that both PKM1 and PKM2 dose-dependently increased the rate of SARM1-catalyzed NAD^+^ cleavage (Fig. 1H). Control experiments using BSA confirmed that NADase potentiation was not an assay artifact (Fig. S1E), and the absence of SIRT1 effects on PKM2 activity demonstrated that the SARM1-PKM interplay is independent of NAD^+^-dependent deacetylation (Fig. S1F). Together, these findings demonstrate that PKM directly binds to SARM1 and allosterically enhances its NADase activity, suggesting a novel regulatory mechanism that links metabolic enzymes to axonal degeneration.

### PKM Potentiates SARM1 NADase via a Phosphorylation-Independent Mechanism

Dimeric PKM2 can act as a protein kinase in non-canonical contexts, raising the possibility that PKM2 phosphorylates SARM1 to drive its activation^18^. To probe this, we generated two PKM2 mutants: R399E, which increases PKM2’s kinase activity^19^, and Q393K, which preserves its pyruvate kinase function but abolishes kinase activity^20^. Both mutants retained robust binding to SARM1 in 293T cells, demonstrating that the PKM2-SARM1 interaction is independent of PKM2’s oligomeric or kinase state (Fig. 2A).

**Figure 2.**
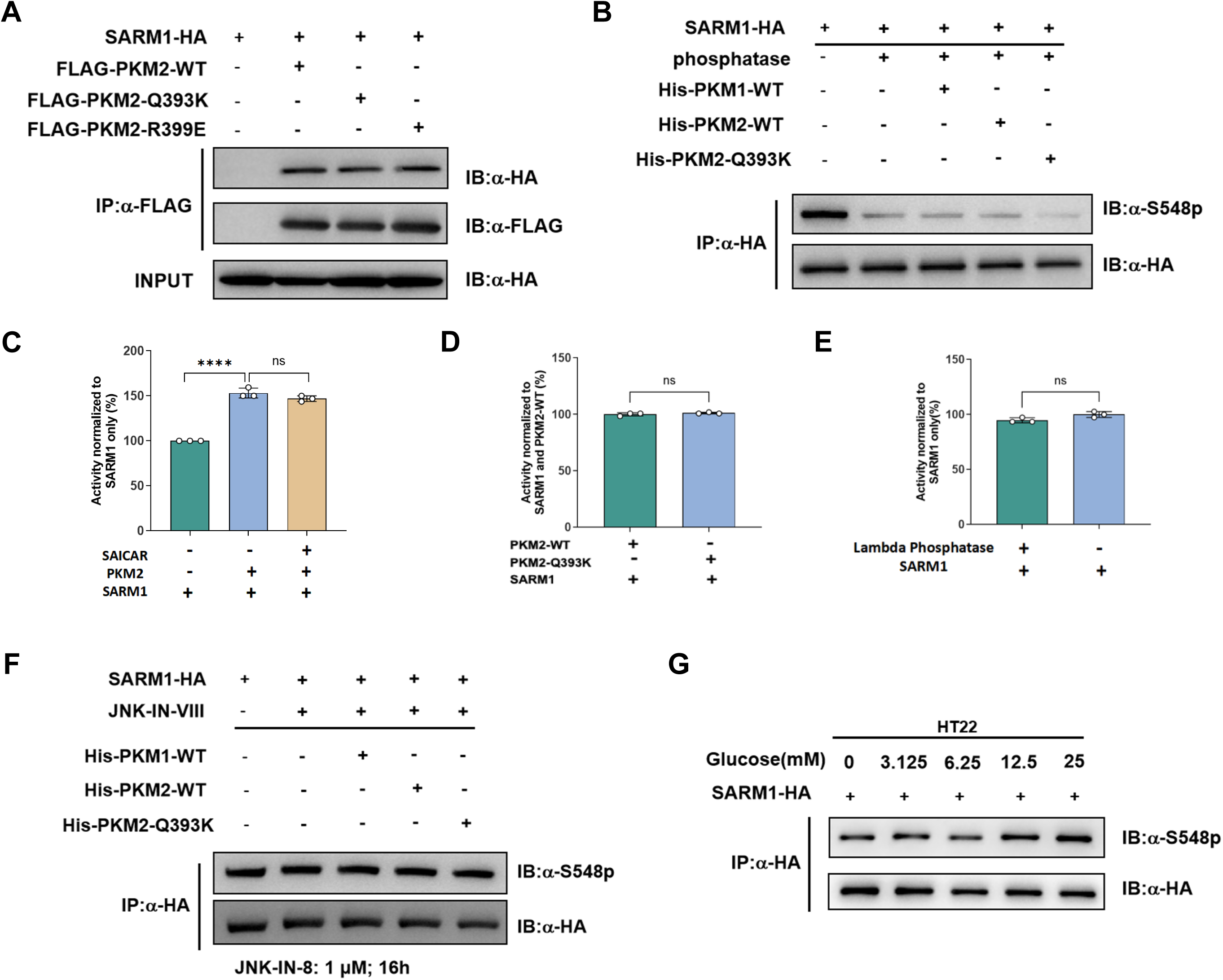
PKM potentiates SARM1 NADase via a phosphorylation-independent mechanism. (**A**) Q393K and R399E mutations of PKM2 do not disrupt its interaction with SARM1. 293T cells were co-transfected with SARM1-HA (2 μg) and either wild-type (WT) FLAG-PKM2 or its mutants (FLAG-PKM2-Q393K/R399E, 2 μg). Cell lysates were immunoprecipitated with anti-FLAG beads and analyzed by western blotting using an anti-HA antibody. Comparable SARM1 binding was observed across WT and mutant PKM2. Input controls (10% total lysate) confirmed equivalent expression of all constructs. (**B**) PKM2 WT and Q393K mutant do not phosphorylate SARM1 at Serine 548. Phosphorylation of SARM1 at S548 was detected by a phospho-S548-specific antibody. PKM1 WT, PKM2 WT, and PKM2-Q393K groups exhibited barely detectable SARM1 phosphorylation compared to the positive control. (**C**) SAICAR does not influence PKM2-mediated enhancement of SARM1 NADase activity. Recombinant SARM1 (6 µg/mL) was incubated with NAD^+^ (2000 μM), NMN (500 µM) and PC6 (50 µM) in the presence or absence of PKM2 and SAICAR. SAICAR had no significant effect on PKM2-dependent SARM1 activation (ns: p > 0.05 vs. PKM2 alone, unpaired t-test). Data are presented as mean ± SD (n = 3). (**D**) PKM2 WT and Q393K mutant equally enhance SARM1 NADase activity. Both PKM2 variants similarly increased the NAD⁺ degradation rate (ns: p > 0.05). Data are presented as mean ± SD (n = 3). (**E**) SARM1 NADase activity is independent of its phosphorylation status. Recombinant SARM1 (6 µg/mL) was treated with λ-phosphatase (100 U, 1 h) or vehicle control. NAD^+^ hydrolysis rates did not significantly differ between groups (ns: p > 0.05, unpaired t-test). Data are presented as mean ± SD (n = 3). (**F**) PKM isoforms do not phosphorylate SARM1 at S548 under JNK inhibition. SARM1-HA was immunoprecipitated from cell lysates using HA beads and incubated with purified PKM1, PKM2, or PKM2-Q393K in the phosphorylation assay following pretreatment with the JNK inhibitor JNK-IN-8 (1 μM, 16 h). Immunoblotting with a phospho-S548-specific antibody detected no increase in phosphorylation levels across all conditions, indicating that PKM does not phosphorylate SARM1 at Ser548. (**G**) SARM1 S548 phosphorylation is insensitive to extracellular glucose levels. HEK293T cells expressing SARM1-HA were cultured in DMEM with glucose (0, 3.125, 6.25, 12.5, 25 mM) for 30 min. Anti-SARM1-S548p immunoblot showed constant phosphorylation levels. Total SARM1-HA expression (anti-HA) confirms equal protein input.

Next, we directly assessed SARM1 phosphorylation at Ser548, a site implicated in its regulation^21^. HA-immunoprecipitated SARM1, treated with λ-phosphatase to remove existing phosphates, was incubated with wild-type PKM1, PKM2, or PKM2-Q393K. Immunoblotting with a phospho-Ser548-specific antibody failed to detect any phosphorylation above background (Fig. 2B). Consistently, addition of SAICAR-a metabolite that enhances PKM2’s kinase activity-did not further augment PKM2-induced SARM1 NADase activation (Fig. 2C). Moreover, both PKM2-WT and the kinase-dead Q393K mutant enhanced SARM1’s NADase rate to the same degree (Fig. 2D), and λ-phosphatase treatment had no appreciable effect on SARM1’s basal or PKM-stimulated NADase activity (Fig. 2E).

To exclude involvement of alternative kinases, we repeated NADase assays in the presence of JNK-IN-8, a selective JNK inhibitor known to block stress-induced SARM1 phosphorylation at Ser548. Neither PKM1 nor PKM2 (WT or Q393K) triggered detectable Ser548 phosphorylation in JNK-inhibited reactions (Fig. 2F), confirming that PKM augments SARM1 activity independently of both its own kinase function and JNK-mediated pathways.

Together, these data establish that PKM activates SARM1 through a phosphorylation-independent mechanism-most likely via direct allosteric modulation-thereby creating a feedforward loop that connects hyperglycemia-driven glycolytic flux to NAD^+^ depletion and axonal degeneration.

### PKM C-Terminal Domain Mediates Direct Binding to the SARM1 SAM-TIR Region

To delineate the molecular interface between PKM and SARM1, we engineered FLAG-tagged PKM truncations corresponding to its N-terminal, A, B and C-terminal domains (CTD)^22^. Co-immunoprecipitation from 293T cells co-expressing SARM1-HA revealed that only the PKM CTD retained robust binding to SARM1, despite lower expression levels of the isolated CTD (Fig. 3A and Fig. S3A). Reciprocally, doxycycline-induced, untagged SARM1 co-precipitated both PKM1-CTD and PKM2-CTD with equivalent efficiency, indicating that the 22-residue differences between isoforms do not affect binding (Fig. 3C). Next, we mapped the corresponding SARM1 interface. GFP-tagged truncations of SARM1-ARM, ARM-SAM, SAM-TIR and full-length were co-expressed with full-length PKM1 in 293T cells. Only the SAM-TIR fragment, which harbors the catalytic TIR domain, showed strong association with PKM1, whereas ARM-SAM and ARM alone failed to co-precipitate PKM1 (Fig. 3B). To test whether the CTD alone could allosterically activate SARM1, we purified recombinant PKM1-CTD (Fig. S3B) and performed in vitro NADase assays. Addition of PKM1-CTD recapitulated the dose-dependent enhancement of SARM1 activity observed with full-length PKM1 (Fig. 3D).

**Figure 3.**
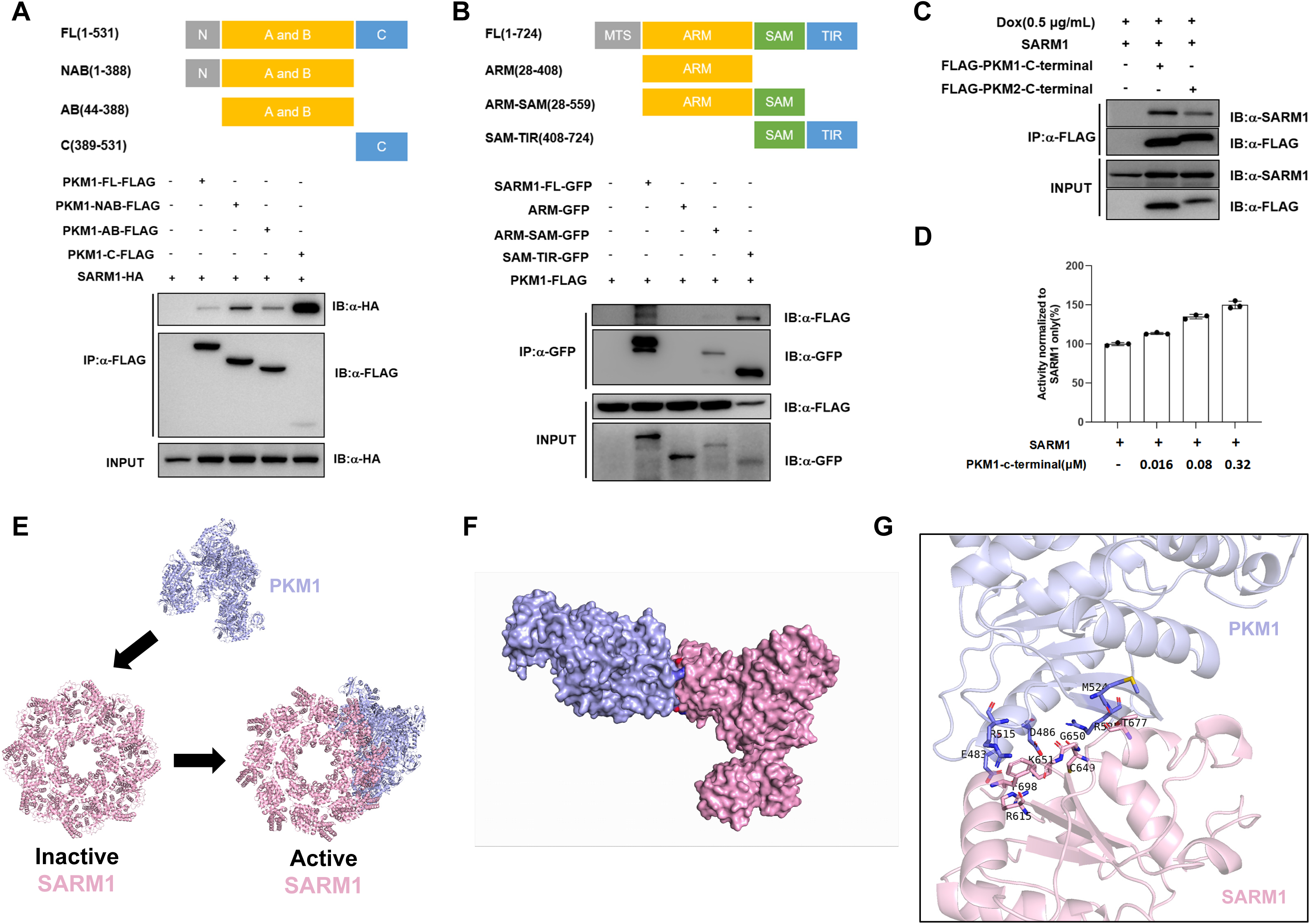
PKM C-terminal domain mediates direct binding to the SARM1 SAM-TIR region. (**A**) The C-terminal domain of PKM1 strongly interacts with SARM1 and enhances its enzymatic activities. 293T cells were co-transfected with full-length SARM1-HA (2 μg) and PKM1-FLAG truncation mutants (2 µg). FL: full-length PKM1 (residues 1-531); NAB: the N-terminal domain, A and B domain of PKM1 (residues 1-388); AB: the A and B domain of PKM1 (residues 44-388); C: the C-terminal domain of PKM1 (residues 389-531). Despite the lower precipitation efficiency of PKM1 C-terminal protein, SARM1 showed the strongest enrichment when co-precipitated with the C-terminal truncation. Input lanes (10% total lysate) showed protein expression levels. (**B**) The TIR domain of SARM1 interacts with PKM1. 293T cells were co-transfected with full-length PKM1-FLAG (2 μg) and SARM1-GFP truncation mutants (2 µg). FL: full-length SARM1 (residues 1-724); ARM: the ARM domain of SARM1 (residues 28-408); ARM-SAM: the ARM and SAM domain of SARM1 (residues 28-559); SAM-TIR: the SAM and TIR domain of PKM1 (residues 408-724). PKM1 showed stronger interaction with SAM-TIR. Input lanes (10% total lysate) showed protein expression levels. (**C**) The C-terminal of PKM1 and PKM2 both interact with SARM1. Co-IP/WB analysis in doxycycline-induced HEK293T cells expressing untagged full-length SARM1 (0.5 μg/mL dox, 16 h) with transient transfection of FLAG-PKM1-CTD or FLAG-PKM2-CTD. Both isoforms strongly co-precipitate endogenous SARM1, despite 22 divergent residues in their C-terminal regions. Input lanes (10% total lysate) showed protein expression levels. (**D**) The C-terminal of PKM1 enhances the enzymatic activities of SARM1. Recombinant SARM1 (6 µg/mL) was incubated with NAD^+^ (2000 μM), NMN (500 µM) and PC6 (50 µM) in the presence or absence of PKM1-CTD (0.016, 0.08, 0.32 µM). Data normalized to control (100% activity = SARM1 without PKM1-CTD). Bars represent standard error. (**E**) Structural modeling suggests that PKM1 induces a conformational change in SARM1, transitioning it from an inactive to an active state. A structural model of the SARM1-PKM1 complex was generated using ZDOCK 3.0.2. The transition from the inactive to the active conformation of SARM1 was observed upon PKM1 binding, indicating an allosteric activation mechanism. (**F**) Crystal structure of SARM1-PKM1 complex predicted by ZDOCK 3.0.2 based on the crystal structure of SARM1 (PDB accession code 7KNQ) and PKM1 (PDB accession code 3SRF). The overall structure of the complex in PyMOL, with SARM1 (light pink) and PKM1 (light blue) shown as surfaces. (**G**) Predicted SARM1 and PKM1 interaction interface. SARM1-TIR interacts with PKM1-C-terminal as shown. The key sites (R615, C649, G650, K651, T677, F698) of SARM1 and the key sites (E483, D486, R515, M524, R525) of PKM1 in the focused panel are shown as sticks and colored in light pink and light blue respectively.

Together, these data establish that the PKM CTD engages the SARM1 SAM-TIR module and is both necessary and sufficient to drive NAD^+^ cleavage, independent of PKM’s canonical glycolytic or kinase functions (Fig. 3E-G), making it a potential therapeutic target. Given the tractability of both domains, these insights pave the way for developing selective inhibitors through rational drug design or peptide-based disruption. Structural modeling of the PKM–SARM1 complex could further guide the development of targeted therapies.

### Hyperglycaemia Enhances SARM1-PKM Complex Formation and Drives NADase Activation

To determine whether elevated glucose promotes SARM1-PKM assembly, we employed our split HaloTag complementation assay in 293T cells. Under increasing glucose concentrations, confocal imaging revealed a marked rise in JF549 positive puncta, indicative of enhanced SARM1-PKM proximity (Fig. 4A). Quantitative analysis of JF549/DAPI fluorescence ratios confirmed a statistically significant, dose-dependent increase in complex formation. Parallel co-immunoprecipitation of endogenous PKM1 with HA-SARM1 likewise showed a glucose-dependent enhancement of binding, despite constant PKM1 expression levels (Fig. 4B). Consistent with our kinase-independent model, SARM1 Ser548 phosphorylation remained unchanged across glucose conditions (Fig. 2G).

**Figure 4.**
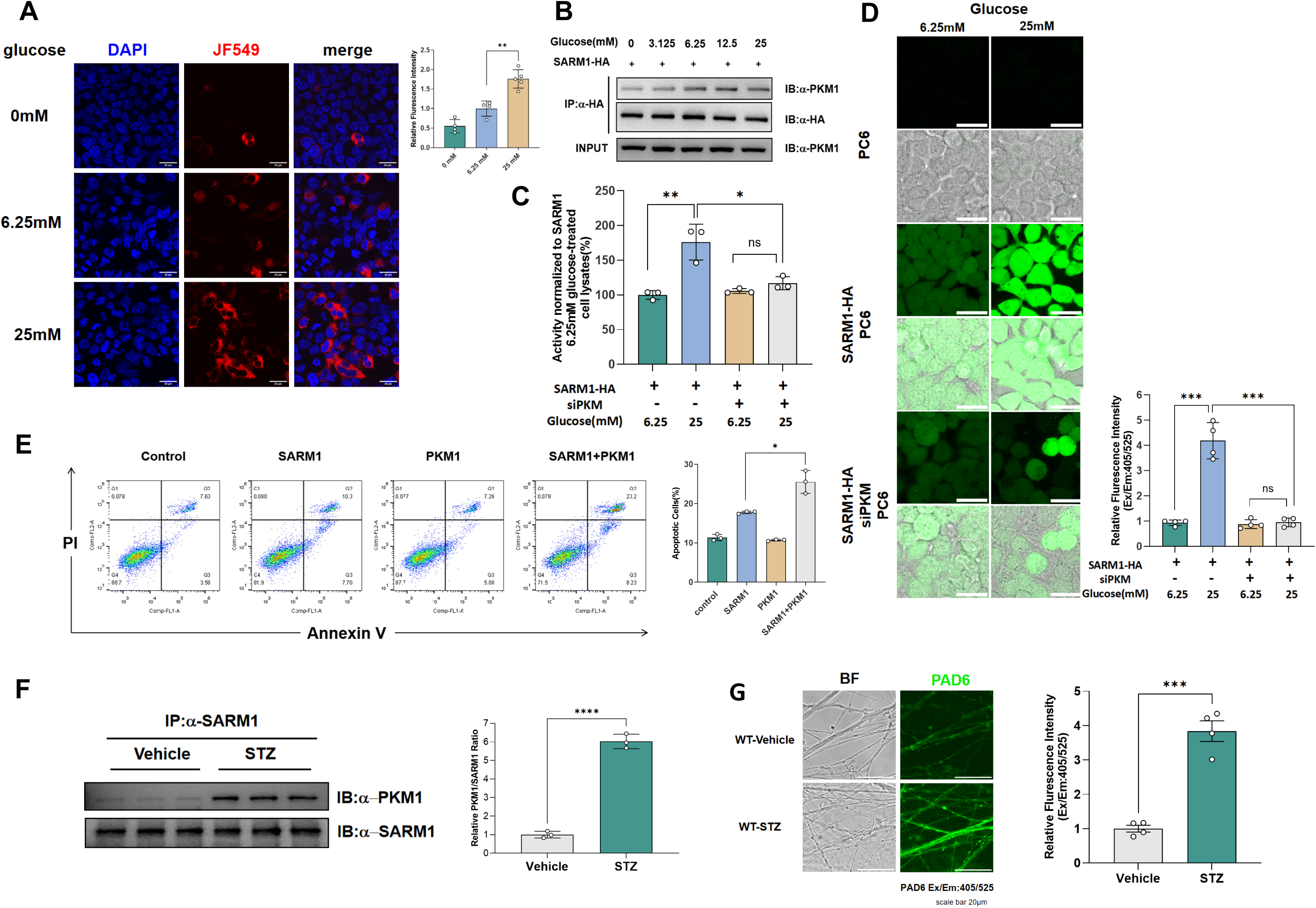
Hyperglycaemia enhances SARM1-PKM complex formation and drives NADase activation. (**A**) High glucose enhances SARM1-PKM interaction via BiFC. HT22 cells co-expressing HaloTag-N58-PKM1-FLAG and SARM1-HaloTag-C58 constructs were cultured in DMEM with glucose (0, 6.25, 25 mM) for 30 min. Red puncta (reconstituted JF549) indicates SARM1-PKM interaction. Nuclei counterstained with DAPI (blue). Right panel: Quantification of relative fluorescence intensity (**p < 0.01 vs. 6.25 mM glucose, unpaired t-test). Scale bar: 20 μm. (**B**) Glucose-dependent SARM1-PKM interaction validated by co-IP. 293T cells over-expressing SARM1-HA were treated with glucose (0, 3.125, 6.25, 12.5, 25 mM) for 30 min. Immunoprecipitation using anti-HA beads followed by probing with anti-PKM1 antibody revealed an increase in PKM1 binding with higher glucose concentrations. Input controls (10% total lysate) confirm comparable PKM1 expression levels across conditions. (**C** and **D**) 293T cells overexpression of SARM1 were pre-transfected with either scramble or PKM-specific siRNA and treated in low (6.25 mM) and high (25 mM) glucose concentrations before the detection of SARM1 activities. (**C**) Cell lysates extracted from the cells were added to a 96-well plate with PC6 assays which contain 50 μM PC6, NADase activities of SARM1 were quantified by fluorescence kinetics of PAD6 from the initial few minutes of reactions. (**D**) Confocal fluorescence images of 293T cells after incubation with 50 μM PC6 and pre-treated with different glucose concentrations, quantification of the cell fluorescence(right). Green: PAD6; scale bar 25 μm. (**E**) Flow cytometric analysis of HT22 cells overexpression of SARM1 or PKM1, and co-overexpression of SARM1 and PKM1, quantification of the cell apoptotic rate (right). (**F**) Co-IP of SARM1 in the cell lysates extracted from DRG neurons of WT C57 mice after 5-day consecutive injection with vehicle or STZ and for the subsequent 6 weeks, quantification of the PKM1/SARM1 ratios (right). (**G**) Confocal imaging of DRG neuronal axons cultured for 7 days, DRG neurons were dissected from WT mice after 5-day consecutive injection with vehicle or STZ and for the subsequent 6 weeks, quantification of the cell fluorescence(right). Green: PAD6; scale bar 20μm. (means ±SD; Student’s t-test,*p<0.05, **p<0.01,****p<0.0001)

We next monitored SARM1 activation using PC6, a selective, cell-permeant probe that is converted by active SARM1 into PAD6, producing a characteristic red-shifted fluorescence. In lysates from 293T cells overexpressing SARM1, PC6 fluorescence kinetics increased significantly with added PKM1 or under high-glucose culture (Fig. 4C). Live cell imaging mirrored these results: hyperglycaemic treatment potentiated PAD6 accumulation in SARM1 expressing cells, an effect abrogated by PKM1 siRNA knockdown (Fig. 4D and Fig. S4A).

Given that axonal damage frequently precedes neuronal injury and death, we assessed the functional consequence of complex formation by measuring apoptosis in HT22 neurons using flow cytometry. SARM1 overexpression alone elevated Annexin V positivity, whereas co-expression with PKM1 further increased the apoptotic fraction. PKM1 overexpression in isolation had no detectable effect (Fig. 4E), underscoring PKM’s role as a facilitator of SARM1 driven neurotoxicity.

Finally, in dorsal root ganglia (DRGs) from streptozotocin-induced diabetic mice, co-IP demonstrated a marked increase in endogenous PKM1-SARM1 interaction compared to controls (Fig. 4F). Similarly, cultured DRG axons incubated with PC6 showed significantly elevated PAD6 fluorescence in diabetic animals, indicating that enhanced SARM1 NADase activity under hyperglycemic conditions (Fig. 4G). Collectively, these findings demonstrate that high glucose promotes PKM-dependent engagement with SARM1, triggering NAD⁺ depletion, axonal fragmentation, and neuronal loss-thereby mechanistically linking diabetic metabolic stress to axon degeneration.

### PKM Knockdown in DRG Protects Against Streptozotocin-induced Neuropathy in vivo

To evaluate the role of PKM in diabetic neuropathy, we delivered AAV9 encoding a PKM-targeted shRNA intraperitoneally into neonatal mice (Fig. 5A). The shRNA was designed to target a sequence common to both PKM1 and PKM2, enabling selective and sustained knockdown of PKM isoforms in dorsal root ganglion (DRG) neurons. Efficient depletion of PKM, without altering SARM1 expression, was confirmed by Western blot and immunohistochemistry (Fig. 5B, C). To determine whether SARM1 mediates the downstream effects of PKM in diabetes-induced axon degeneration, we included SARM1 knockout (KO) mice in the streptozotocin (STZ)-induced diabetic neuropathy model^23^. We first assessed metabolic baseline, nociceptive thresholds, tactile sensitivity, and motor coordination across all four groups^24^. Neither PKM knockdown nor SARM1 deletion significantly impacted these parameters under STZ treatment (Fig. S3C-J).

**Figure 5.**
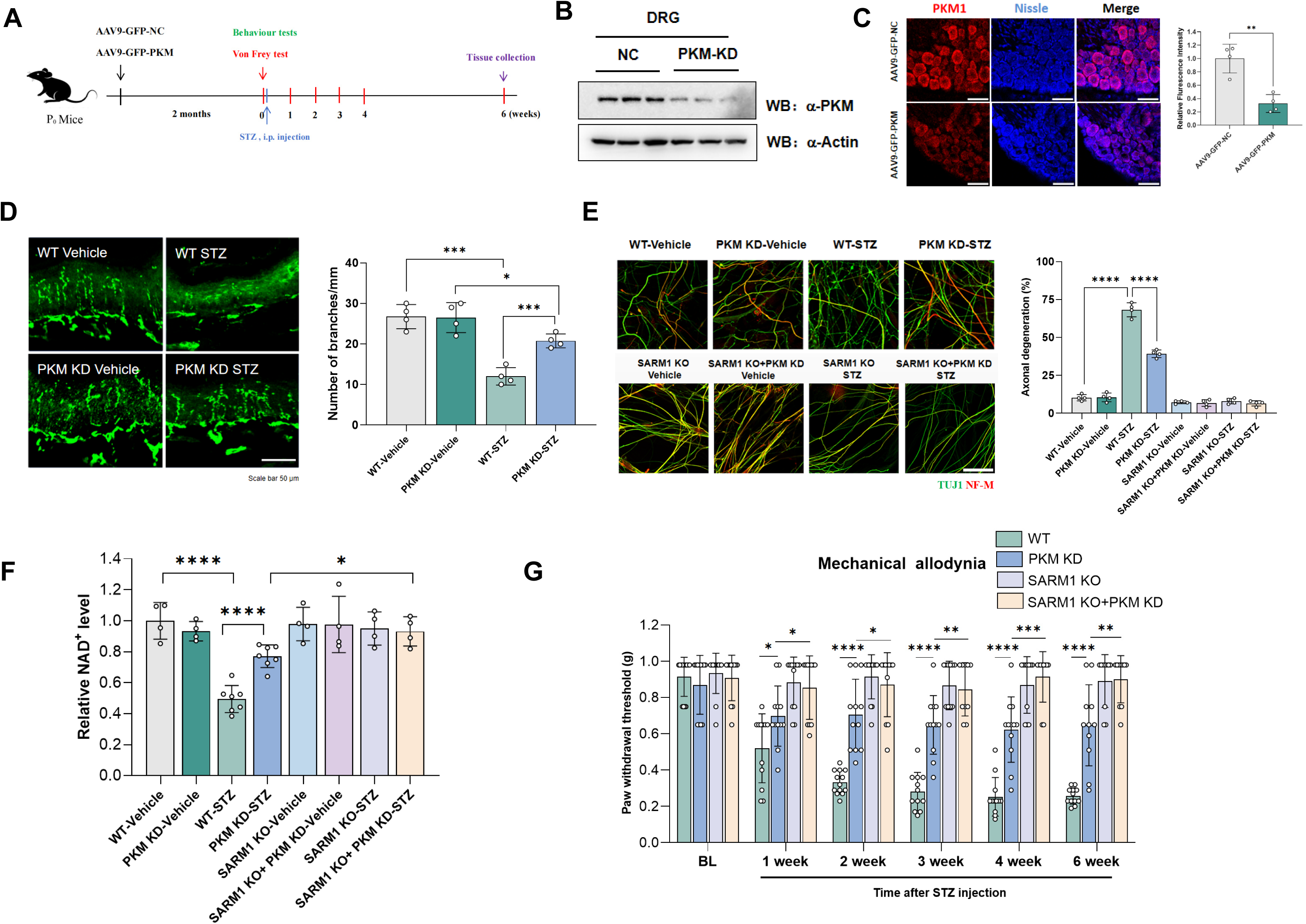
PKM knockdown in DRG protects against streptozotocin-induced neuropathy in vivo. (**A**) Schematic showing the timeline of PKM KD mice preparation, STZ treatment, behavioral tests, and tissue collection from neonatal mice. i.p., intraperitoneal. (**B**) Knockdown efficiency of PKM KD tested by western blotting. NC: AAV9-GFP-NC; PKM KD: AAV9-GFP-PKM. (**C**) Knockdown efficiency of PKM KD tested by immunohistochemistry of frozen sections, quantification of the fluorescence intensity (right). (**D**) Representative images of immunostaining showing intraepidermal nerve fiber in the glabrous skin of hind paw from WT and PKM KD mice after 5-day injection with vehicle or STZ and for the subsequent 6 weeks. Quantification of the density of intraepidermal PGP9.5^+^ nerve fiber branches (right). N = 5 mice per group; scale bar: 50 μm. **(E)** The DRG neuronal axonal degeneration caused by the STZ-induced diabetic neuropathy in WT, PKM KD, SARM1 KO and SARM1 KO and PKM KD mice. The axonal integrity was examined by anti-TUJ1 (green) / anti-NF-M (red) co-immunostaining. Quantification of the axonal degeneration (right). n = 5 mice per group; scale bar: 50 μm. **(F)** The NAD^+^ levels of DRGs in WT, PKM KD, SARM1 KO and SARM1 KO mice after 5-day consecutive injection with vehicle or STZ and for the subsequent 6 weeks were measured. n = 4 to 6 mice per group. (means ± SD; Student’s t-test, *p<0.05, **p<0.01, ****p<0.0001) (**G**) Time course of STZ-induced mechanical allodynia in WT, PKM KD, SARM1 KO and SARM1 KO and PKM KD mice. n = 9 to 12 mice per group.

Intraepidermal nerve fibers (IENFs) in hindpaw skin serve as a key marker of peripheral neuropathy in mice^25^. STZ-treated wild-type animals showed a significant reduction in IENF density, whereas PKM knockdown partially preserved these fibers, and SARM1 knockout mice were fully protected from fiber loss (Fig. 5D). To further examine DRG neuron integrity, we cultured DRG neurons from diabetic mice. Axonal fragmentation was prominent in STZ-treated wild-type neurons, but markedly reduced in PKM-knockdown mice and entirely absent in SARM1-KO neurons (Fig. 5E). Consistent with these observations, STZ-induced NAD⁺ depletion in DRGs was significantly mitigated by PKM knockdown, and entirely absent in SARM1-KO ganglia (Fig. 5F and Fig.S5C). Behavioral assessment using the von Frey assay revealed robust mechanical allodynia in diabetic wild-type mice. PKM-knockdown mice retained significantly higher withdrawal thresholds, and SARM1-KO mice were fully protected from allodynia (Fig. 5G).

Together, these findings demonstrate that PKM is essential for SARM1-mediated NAD^+^ depletion, axonal degeneration, and pain hypersensitivity in diabetic peripheral neuropathy, and highlight the therapeutic potential of targeting the PKM-SARM1 axis for neuroprotection in metabolic neurodegeneration.

### Pharmacological Disruption of the SARM1-PKM Interface Ameliorates Diabetic Neuropathy

Building on Figure 3, which demonstrated that the PKM C-terminal domain engages the SARM1 SAM-TIR module-highlighting a structurally tractable interface for selective inhibition-we sought to therapeutically disrupt this interaction. Using AlphaFold-based structural modeling, we designed a competitive inhibitory peptide (Pep-SP1) derived from residues 645–655 of the SARM1 TIR domain (Fig. 6A). In AlphaScreen binding assays, Pep-SP1 effectively inhibited the SARM1-PKM1 interaction in a dose-dependent manner, yielding an IC_50_ of 1.068 µM (Fig. 6B). In HEK293T cells co-expressing FLAG-PKM1 and SARM1-HA, treatment with Pep-SP1 (1 or 10 µM) significantly reduced co-immunoprecipitation of SARM1 with PKM1, relative to a scrambled control peptide (Fig. 6C). This inhibitory effect was further corroborated by in vitro pull-down assays using recombinant FLAG-SARM1 and His-PKM1, confirming the ability of Pep-SP1 to block the interaction between these proteins (Fig. 6D).

**Figure 6.**
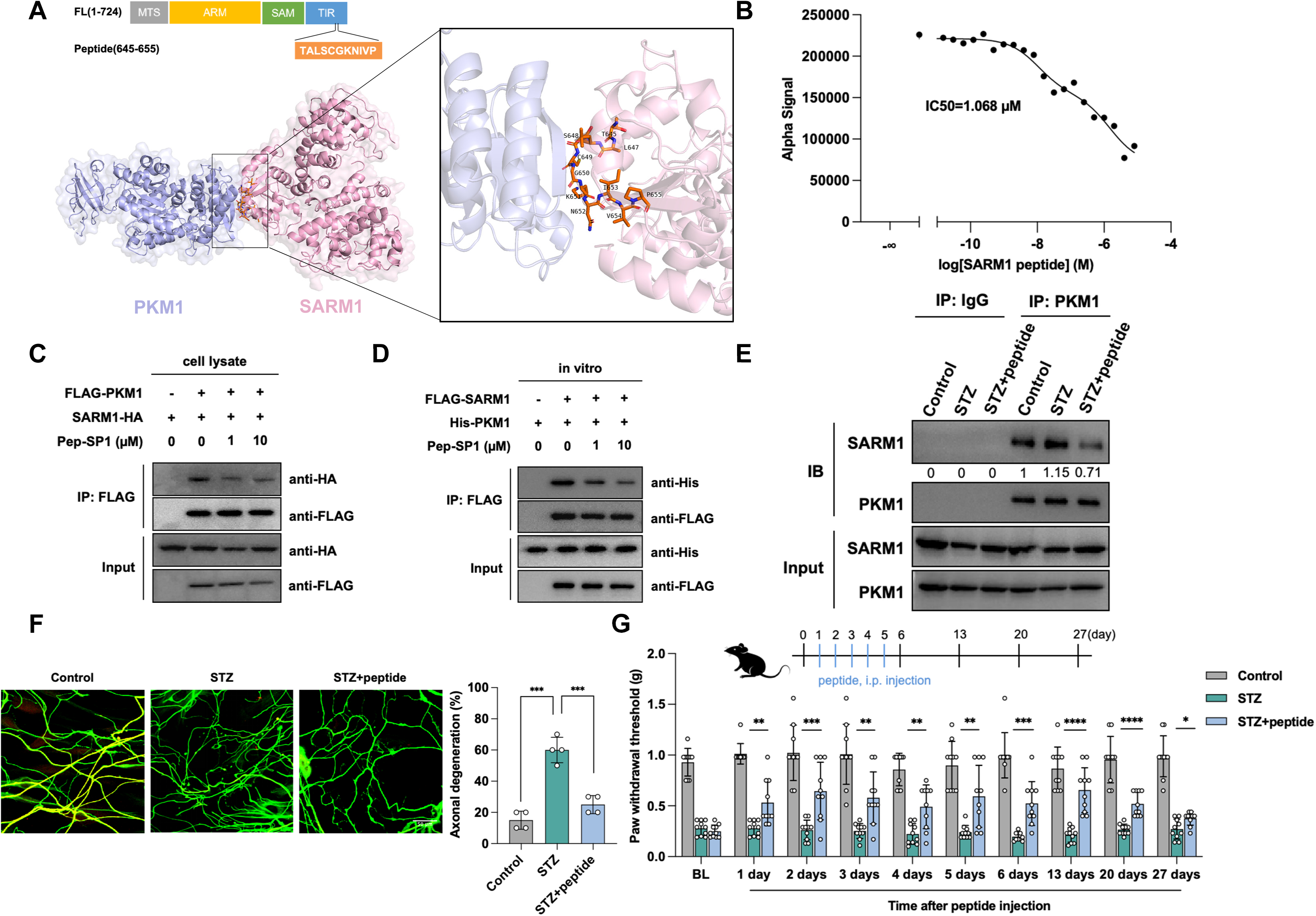
Pharmacological disruption of the SARM1–PKM1 interface ameliorates diabetic neuropathy. (**A**) Structural modeling of Pep-SP1 bound to the SARM1-PKM1 complex. The peptide (orange) occupies the SARM1-PKM1 interaction interface, disrupting their binding. (**B**) Dose-dependent inhibition of SARM1-PKM1 binding by Pep-SP1, as determined by AlphaScreen assay (IC50 = 1.068 µM). (**C**) Pep-SP1 reduced SARM1-PKM1 interaction compared to scrambled peptide controls. SARM1-HA (2 μg) and FLAG-PKM1 (2 μg) plasmids were co-expressed for 24 h in 293T cells. Lysates were treated with Pep-SP1 (1, 10 µM) for 3 h and immunoprecipitated with anti-FLAG antibody. Input controls showed transfection efficiency. (**D**) In vitro co-IP assay with recombinant FLAG-SARM1 and His-PKM1. Pep-SP1 (1, 10 µM) dose-dependently disrupted SARM1-PKM1 binding. (**E**) Co-IP analysis of DRG lysates from STZ-induced diabetic mice. SARM1-PKM1 interaction increased by 1.15-fold in diabetic mice (vs. non-diabetic controls) and was reduced to 0.71-fold after daily Pep-SP1 treatment (10 mg/kg). (**F**) Axonal integrity in DRG neurons. Control: normal mice; STZ: STZ-induced diabetic mice; STZ+peptide: peptide-injected STZ-induced diabetic mice. STZ-induced diabetic mice exhibited increased axonal fragmentation (fragmentation index: 0.6 ± 0.08 vs. non-diabetic), which was significantly mitigated by Pep-SP1 (0.25 ± 0.05, ***p < 0.001). Representative immunofluorescence images are shown. n = 3 mice per group; scale bar: 50 μm. (**G**) Time course of mechanical allodynia. STZ-induced diabetic mice displayed hypersensitivity, partially reversed by Pep-SP1 (thresholds: 0.66 ± 0.2 g at day 13 vs. 0.38 ± 0.07 g at day 27), suggesting transient efficacy. n = 10 mice per group. (means ± SD; Student’s t-test, *p<0.05, **p<0.01, ***p<0.001, ****p<0.0001)

We next evaluated the in vivo efficacy of Pep-SP1 in a STZ-induced mouse model of diabetic peripheral neuropathy. Co-immunoprecipitation of DRG lysates revealed that a 1.15-fold increase in the endogenous SARM1–PKM1 interaction in diabetic mice compared to non-diabetic controls. Notably, daily intraperitoneal administration of Pep-SP1 (10 mg/kg) over four weeks significantly reduced this interaction to 0.71-fold of baseline levels (Fig. 6E). Importantly, this treatment regimen was well tolerated, with no detectable hepatic or renal toxicity (Fig. S6G, H).

Functionally, Pep-SP1 treatment markedly reduced axonal degeneration in DRG neurons, as indicated by a significant decrease in the axonal fragmentation index (0.25 ± 0.05 vs. 0.60 ± 0.08 in vehicle-treated STZ mice; P < 0.001; Fig. 6F and Fig.S6L). Moreover, Pep-SP1 alleviated mechanical allodynia in diabetic mice, with paw withdrawal thresholds improving progressively to 0.66 ± 0.2 g by day 13 post-treatment. However, this effect diminished by day 27 (0.38 ± 0.07 g), potentially reflecting limited in vivo peptide stability or the activation of compensatory neurodegenerative pathways (Fig. 6G).

Together, these findings demonstrate that targeted disruption of the SARM1-PKM interface suppresses SARM1 activation, preserves axonal integrity, and mitigates sensory deficits in diabetic neuropathy, underscoring its potential as a therapeutic strategy for metabolic neurodegeneration.

## Discussion

Diabetic peripheral neuropathy (DPN) remains a prevalent and disabling complication of diabetes^2627^, yet its molecular underpinnings remain incompletely understood, and effective treatments are lacking. Here, we identify a previously unrecognized role for pyruvate kinase M (PKM) as a glucose-sensitive activator of the axonal degeneration mediator SARM1, linking metabolic dysregulation to neurodegeneration. We show that hyperglycemia promotes the formation of a SARM-PKM complex, which in turn amplifies SARM1’s NADase activity and drives axonal loss^28^. This interaction is structurally mediated by the C-terminal domain of PKM1 and the TIR domain of SARM1 and is independent of PKM1’s canonical enzymatic function in glycolysis^29^. Disruption of this interface with a rationally designed peptide attenuates neuropathological features in vivo, establishing the SARM1-PKM axis as both a mechanistic driver and a druggable target in DPN.

These findings provide mechanistic insight into how metabolic perturbations, particularly hyperglycemia, activate SARM1 in sensory neurons. While SARM1 activation and subsequent NAD^+^ depletion have been implicated in axon degeneration across models^30^, the upstream metabolic cues regulating this process in diabetic conditions have remained unclear. Our results position PKM as a critical node that senses glucose availability and, through direct interaction with SARM1, relays metabolic stress into a degenerative signal. The kinase-independent activation of SARM1 by PKM expands the functional repertoire of glycolytic enzymes in neurons^31^, suggesting that metabolic enzymes can moonlight as direct effectors of neurodegenerative pathways.

Our data support a model in which hyperglycemia induces a self-reinforcing feedforward loop: PKM binding enhances SARM1 activation, which in turn accelerates NAD^+^ depletion, mitochondrial dysfunction, and axonal degeneration-hallmarks of DPN pathology^32^. Despite the 22-residue difference in their C-terminal domains and their distinct tissue distributions and regulatory properties, both PKM1 and PKM2 exhibited similar capacities to activate SARM1 under hyperglycemic conditions in our study. Genetic silencing of PKM in dorsal root ganglia (DRG) preserved intraepidermal nerve fiber density, maintained NAD^+^ levels, and mitigated mechanical allodynia in diabetic mice, consistent with disruption of this loop. Moreover, the peptide Pep-SP1, designed to competitively block the SARM1–PKM interaction, conferred neuroprotection in vivo, providing proof-of-concept for therapeutic intervention. This work raises broader questions regarding the role of PKM–SARM1 crosstalk in other neurodegenerative conditions associated with metabolic stress. Altered expression of PKM isoforms has been reported in Alzheimer’s and Parkinson’s disease^333435^, and SARM1 is increasingly recognized as a conserved mediator of axon loss across diverse contexts^73637^. The structural and functional conservation of the SARM1–PKM interaction suggests that this axis may represent a shared mechanism linking metabolic imbalance to neuronal injury beyond diabetic neuropathy.

Recent studies have uncovered multiple strategies to inhibit SARM1, including orthosteric inhibition via enzyme-catalyzed base-exchange reactions and allosteric blockade of NMN binding, which prevents activation-induced conformational changes^38394041^. Our findings introduce a complementary therapeutic avenue by targeting a previously unrecognized regulatory interface between SARM1 and PKM. The TIR-derived peptide inhibitor (Pep-SP1) we developed selectively disrupts this interaction, thereby attenuating SARM1 activation in response to hyperglycemic stress.

This approach circumvents the need to directly block the catalytic or allosteric sites of SARM1, instead acting upstream by preventing activator-induced NADase engagement. Given the structural modularity of the PKM-SARM1 interface, this strategy may afford improved specificity and reduced off-target effects compared to active site-directed inhibitors. Moreover, as protein–protein interaction interfaces often mediate context-dependent activation, this paradigm may enable disease-selective modulation of SARM1 without compromising its basal physiological functions. Thus, our work expands the repertoire of SARM1-targeted interventions and highlights the therapeutic potential of interface-based inhibition in neurodegenerative conditions driven by metabolic dysregulation.

In summary, we define a glucose-sensitive SARM1-PKM axis as a key effector of axonal degeneration in DPN, revealing a direct molecular link between metabolic dysregulation and neuronal damage. By targeting the interface between a glycolytic enzyme and a pro-degenerative NADase, we propose a therapeutic strategy that may extend beyond DPN to a broader spectrum of neurodegenerative diseases at the intersection of metabolism and neuronal survival^42^.

## Acknowledgments

We thank all members of the Yu Laboratory for their valuable technical assistance and insightful discussions. We are grateful to the Facility Center of the State Key Laboratory of Genetics and Development of Complex Phenotypes, particularly the Flow Cytometry Core, for technical support. We thank Prof. Qiwei Zhai for generously providing the SARM1 knockout mice, Prof. Qingjian Han for assistance with behavioral assays and Prof. Yujie Sun for kindly providing the plasmids of Bimolecular fluorescence detection TagBiFC. This work was supported by the National Key R&D Program of China (2023YFA1800400) and the National Natural Science Foundation of China (grant numbers 32370825, 92249302, and 92049301).

## Author contributions

W.Y. conceived and supervised the project, analyzed the data, and wrote the manuscript with input from J.L. and W.L. W.L. and C.L. performed molecular and cellular experiments, including co-immunoprecipitation, NADase activity assays, and live-cell imaging with the help from Z.Z., Y.J., W.Z. and Y.Z. J.L. conducted structural interaction mapping and in vitro binding assays. J.L. and W.L. carried out animal studies, including diabetic mouse models, behavioral assessments, dorsal root ganglion (DRG) neuron cultures, and axonal degeneration assays. J.L. also designed the SARM1-PKM inhibitory peptide and validated its therapeutic efficacy in vivo. All authors contributed to data interpretation and manuscript revision.

## Competing interests

The authors declare no competing interests.

## Methods

### Antibodies and reagents

The following primary antibodies were used in this study: anti-FLAG (Sigma-Aldrich, SAB4301135; 1:4,000), anti-HA (Abcam, ab9110; 1:4,000), anti-GFP (Abcam, ab290; 1:1,000), anti-His (ABclonal, AE028; 1:4,000), anti-SARM1 (Cell Signaling Technology, 13022; 1:1,000), anti-PKM1 (Cell Signaling Technology, 7067; 1:1,000), and anti-PKM2 (Cell Signaling Technology, 4053; 1:1,000). Biochemical reagents including NAD⁺, NMN, TEMED, N, N′-Methylenebisacrylamide, phenylmethanesulfonyl fluoride (PMSF), aprotinin, leupeptin, pepstatin A, sodium orthovanadate (Na₃VO₄), sodium fluoride (NaF), nicotinamide (NAM), trichostatin A (TSA), and DNase-I were obtained from Sigma-Aldrich. General laboratory reagents such as bovine serum albumin (BSA), Tris, HEPES, NaCl, glycine, glycerol, Tween-20, kanamycin, ampicillin, and isopropyl β-D-thiogalactopyranoside (IPTG) were purchased from BBI Life Sciences.

### Plasmid construction

Full-length human SARM1 and PKM1/PKM2 cDNAs were amplified by PCR and subcloned into the following expression vectors: pRK7-C-HA, pcDNA3.1-N-FLAG, pET28a-His-SUMO, and pCDH (Addgene plasmid #64875). HaloTag-N58-c-Jun and HaloTag-c-Fos-C58 were generously gifted by Prof. Yujie Sun (Peking University). Transcriptional coactivators Tip60, PCAF, and GCN5 were cloned into the pcDNA3.1-HA vector. Truncated SARM1 domain constructs including ARM-GFP, ARM-SAM-GFP, and SAM-TIR-GFP were generously provided by Dr. Zhe Zhang (Peking University). All constructs were verified by Sanger sequencing before use in cellular or biochemical assays.

### Cell lines and transfection

HEK293T and HT22 cells were cultured in Dulbecco’s Modified Eagle’s Medium (DMEM, Gibco) supplemented with 10% (v/v) fetal bovine serum (FBS) and 0.1 mg/mL streptomycin and 100 U/mL penicillin at 37 ℃ with 5% CO2. Expi293F cells (a gift from Prof Lin Jinzhong, Fudan University) were cultured at 37 °C with 5% CO2 in serum-free SMM-293TII (Sino Biological) medium. Transient transfections were performed using Lipofectamine 2000 (Invitrogen) according to the manufacturer’s protocol. Cells were transfected at approximately 60% confluence and harvested 24– 48 hours post-transfection for downstream analyses.

### Expression and purification of PKM Proteins

The full-length human PKM1 cDNA was cloned into the pET28a vector containing an N-terminal His-SUMO tag. Recombinant PKM1 protein was expressed in Escherichia coli BL21 (DE3) cells. Briefly, bacterial cultures were grown overnight in LB medium containing 50 μg/mL kanamycin at 37 °C, then diluted 1:100 into 2×YT medium for large-scale expression. Cultures were incubated at 37 °C until reaching an OD₆₀₀ of 0.6– 0.8, induced with 0.1 mM isopropyl β-D-1-thiogalactopyranoside (IPTG), and incubated overnight at 18 °C. Cells were harvested by centrifugation, washed with PBS, and stored at −80 °C until use. For purification, cell pellets were resuspended in lysis buffer (50 mM HEPES pH 8.0, 500 mM NaCl, 5% glycerol, 20 mM imidazole, and 100 μM phenylmethylsulfonyl fluoride [PMSF]) and lysed using a low-temperature high-pressure homogenizer (JN-3000 Plus). The lysates were clarified by centrifugation at 18,000 rpm for 1 h at 4 °C. The supernatant was loaded onto a 5 mL HisTrap FF Ni-Sepharose column (GE Healthcare) and eluted using a linear imidazole gradient on an ÄKTA FPLC system (GE Healthcare). The elution buffer consisted of Buffer A (same as lysis buffer) and Buffer B (identical to Buffer A except containing 500 mM imidazole). The His-SUMO tag was removed by overnight digestion with ULP1 protease at 4 °C. Recombinant PKM2, the PKM2-Q393K mutant and the C-terminal domain of PKM1 were expressed and purified following the same protocol.

### Expression and purification of SARM1

The human SARM1 cDNA, fused at the N-terminus with a twin-Strep-tag and a Flag-tag followed by a TEV protease cleavage site, was cloned into the pCDH vector (Addgene plasmid #64875). Recombinant SARM1 was expressed in Expi293F cells cultured in serum-free medium (SMM-293TII, Sino Biological) under standard conditions (37 °C, 5% CO₂, 120 rpm). Transfection was performed using polyethyleneimine (PEI) at a ratio of 1 mg DNA per liter of culture when cell density reached 1 × 10⁶ cells/mL. Cells were harvested 4 days post-transfection by centrifugation at 5,000 rpm for 10 min at 4 °C and resuspended in lysis buffer (100 mM Tris-HCl pH 8.0, 150 mM NaCl, 1 mM EDTA) supplemented with protease inhibitors (1 μg/mL aprotinin, leupeptin, and pepstatin; 1 mM PMSF) and DNase I (2 μg/mL). Cells were lysed using a low-temperature high-pressure cell disrupter (JN-3000 Plus), and lysates were clarified by centrifugation at 18,000 rpm for 1 h at 4 °C. The supernatant was filtered through a 0.45 μm membrane and applied to a pre-equilibrated Strep-Tactin affinity column (IBA Lifesciences). After extensive washing with lysis buffer, bound proteins were eluted with buffer containing 2.5 mM D-desthiobiotin. The eluted SARM1 protein was subjected to overnight TEV protease digestion at 4 °C to remove the affinity tags. The resulting protein was either used immediately or further purified as needed.

### Surface plasmon resonance (SPR) analysis

Surface plasmon resonance (SPR) experiments were performed using a Biacore T200 system (GE Healthcare) to assess the binding affinity between SARM1 and PKM1/2. Recombinant SARM1 was diluted in 10 mM sodium acetate buffer (pH 4.0) and immobilized onto a CM5 sensor chip (GE Healthcare) via standard amine coupling chemistry, achieving a final surface density of approximately 4000 response units (RU). Binding interactions were evaluated in single-cycle mode. Increasing concentrations of PKM1 or PKM2 (0, 0.00977, 0.0195, 0.039, 0.0781, 0.1563, 0.3125, 0.625, 1.25, 2.5, 5, 10, and 20 μM) were injected over the immobilized SARM1 at a flow rate of 30 μL/min for 120 s. Following the association phase, dissociation was monitored by injecting running buffer alone (HBS-T supplemented with 5% DMSO) at the same flow rate for an additional 120 s. All measurements were conducted at 25 °C. Sensorgrams were reference-subtracted and analyzed using Biacore T200 Evaluation Software (GE Healthcare) to calculate kinetic parameters and binding affinities.

### *In vitro* SARM1 activity assay

SARM1 enzymatic activity was measured using a fluorescent NADase activity assay. Reactions were initiated by incubating purified SARM1 protein with a reaction mixture containing 50 μM PC6, 2 mM NAD⁺, and 500 μM NMN in 10 mM phosphate-buffered saline (PBS, pH 7.4). Fluorescence was recorded in black 96-well plates using an Infinite M200 PRO microplate reader (Tecan). The reaction kinetics were monitored at an excitation wavelength of 390 nm and an emission wavelength of 520 nm. The initial rate of fluorescence increase, reflecting NADase activity, was quantified by calculating the slope over the linear phase of the reaction.

### Cellular SARM1 activity assay

To assess SARM1 activity in cells, HEK293 cells were seeded on poly-L-lysine-coated chambered coverglass (Thermo Fisher, #155411) and cultured overnight. SARM1 was overexpressed in cells with or without PKM knockdown via siRNA. Cells were then incubated in media containing various glucose concentrations, followed by treatment with 50 μM PC6. For primary neuron studies, dorsal root ganglion (DRG) neurons were isolated from diabetic and healthy mice and cultured on poly-L-lysine-coated chambered coverglass. After 24 hours, neurons were treated with 50 μM PC6. Fluorescence signals corresponding to PAD6 activity (Ex/Em: 405/525 nm) were captured using a Nikon A1 confocal microscope. Images were analyzed to assess SARM1 activation in different conditions.

### PKM Knockdown and Quantitative Reverse Transcription PCR (qRT-PCR)

For gene silencing, 40 pM siRNAs targeting PKM (synthesized by GenePharma) were transfected into cells using Lipofectamine RNAiMAX Reagent (Invitrogen), according to the manufacturer’s protocol. Cells were incubated for 24–48 hours post-transfection before harvesting. Total RNA was extracted using TRIzol Reagent (Invitrogen) following standard procedures. Reverse transcription was performed using ABScript III RT Master Mix with gDNA remover (ABclonal, RK202429). Quantitative PCR was conducted using SYBR Green-based detection, and gene expression levels were normalized to internal control genes. Relative expression was calculated using the ΔΔCt method.

### Flow cytometry

Apoptosis analysis was performed using the Annexin V-FITC/PI Apoptosis Detection Kit (Yeasen, #40302ES60) following the manufacturer’s instructions. Briefly, 5 × 10⁵ cells were harvested using EDTA-free trypsin, centrifuged at 300 × g for 5 min at 4 °C, and washed twice with cold PBS. After discarding the supernatant, cells were resuspended in 100 μL of 1× Binding Buffer. Then, 5 μL of Annexin V-FITC and 10 μL of propidium iodide (PI) staining solution were added to the suspension, mixed gently, and incubated at room temperature for 15 min in the dark. After staining, 400 μL of 1× Binding Buffer was added, and samples were analyzed within 1 hour using a FACSVerse flow cytometer (BD Biosciences). Data were processed and analyzed using FlowJo software (Tree Star).

### Animals

All animal procedures were conducted under the guidelines set forth by the International Association for the Study of Pain and were approved by the Animal Care and Use Committee of Fudan University. Male C57BL/6 mice were purchased from the Institute of Developmental Biology and Molecular Medicine at Fudan University. SARM1 knockout (Sarm1^-/-^) mice, with a disruption of exons 3–6, were provided by Dr. Zhai Qiwei from the Shanghai Institute of Nutrition and Health, Chinese Academy of Sciences. Mice were group-housed in cages with 2–5 animals per cage, maintained on a 12-hour light/12-hour dark cycle, and had ad libitum access to food and water. Animals were randomly assigned to different experimental groups.

### Diabetes models

Ten-week-old male C57BL/6 and Sarm1^-/-^ mice were induced with diabetes by intraperitoneal injection of 50 mg/kg/day streptozotocin (STZ, Sigma-Aldrich) dissolved in sodium citrate buffer (0.1 M, pH 4.5). Control mice received an intraperitoneal injection of an equivalent volume of sodium citrate buffer. STZ injections were administered for 5 consecutive days as previously described. Mice with blood glucose levels exceeding 250 mg/dL for two consecutive weeks were considered diabetic.

### Neonatal intraperitoneal (i.p.) injection of AAV

The scAAV9 containing shRNA targeting *pkm* was purchased from WZ Biosciences (China) and administered via neonatal intraperitoneal (i.p.) injection. To facilitate injection, urine was gently expelled from the bladder of male neonatal mice by applying mild abdominal pressure. A total volume of 10 μL of AAV9, with a titer of 4×10¹² vg/mL, was injected intraperitoneally using a Hamilton microsyringe. Following the injection, the injection site was gently pressed for 2 minutes to prevent viral leakage. This method resulted in highly selective AAV9 infection in ganglionic neurons, with minimal infection observed in skin cells and cortical neurons.

### Primary cultures of mouse DRG neurons

Dorsal root ganglia (DRGs) were collected from male wild-type (WT) or Sarm1^-/-^ mice for primary neuronal cultures. Freshly isolated DRGs were immediately digested with an enzyme mixture consisting of deoxyribonuclease I (0.1 mg/mL, Sigma, DN25), trypsin (0.4 mg/mL, Sigma, T8003), and collagenase type IA (1 mg/mL, Sigma, C9891) at 37°C for 45 minutes. After digestion, the neurons were resuspended in Neurobasal Medium (Gibco, LOT2660082), and the cell suspension was passed through a 100-μm cell filter to obtain a single-cell suspension. These DRG cells were plated on glass coverslips pre-coated with poly-L-lysine and cultured in Neurobasal Medium supplemented with 2% B27 supplement (Invitrogen), 10% (v/v) fetal bovine serum (FBS), 0.1 mg/mL streptomycin, and 100 U/mL penicillin at 37°C with 5% CO2. To prevent non-neuronal cell growth, a mitotic inhibitor mixture (5 μM 5-fluoro-2’-deoxyuridine and 5 μM uridine) was added to the medium. The neurons were cultured in a fresh Neurobasal Medium every 48 hours. After 7 days in culture, axonal growth was assessed under a phase-contrast microscope. In some cases, axons were co-immunostained using mouse anti-TUJ1 (1:1,000 dilution, BioLegend, Cat:801201) and chicken anti-NF-M (1:1,000 dilution, BioLegend, Cat:822701), followed by Alexa Fluor 594-conjugated Donkey Anti-Mouse IgG(H+L) (YEASEN, Cat:34112ES60) and Alexa Fluor 647-conjugated Rabbit Anti-Chicken IgY(H+L) (YEASEN, Cat:34713ES60). Immunostained axons were visualized using a confocal microscope (Nikon A1). Axon degeneration was quantified by calculating the degeneration index, defined as the ratio of degenerated axon area (e.g., blebbing or fragmentation) to the total axon area. Four replicate wells were used for each experimental condition.

### Immunohistochemistry

Mice were deeply anesthetized with 1.25% Tribromoethanol (LAT) at a dosage of 0.2 mL/10g body weight and transcardially perfused with 0.01M PBS followed by 4% paraformaldehyde (PFA). After perfusion, the dorsal root ganglia (DRGs) and hind paw glabrous skin were carefully removed. The tissues were post-fixed in 4% PFA overnight, followed by transfer to 30% sucrose at 4°C for dehydration. DRG sections (10 μm) and skin sections (30 μm) were prepared for immunofluorescence staining. Sections were blocked with 2% bovine serum albumin (BSA) dissolved in 0.01M PBS for 1 hour at room temperature. Subsequently, sections were incubated with primary antibodies overnight at 4°C, followed by incubation with secondary antibodies at room temperature for 2 hours. The stained sections were examined under a Nikon fluorescence microscope. At least three sections from each animal were analyzed, with a total of three animals per group. Intraepidermal nerve fiber density (INFD) was assessed according to a previously modified protocol. Nerve fiber profiles were counted blindly, and the INFD was expressed as the mean number of fibers per linear millimeter of epidermis from four sections per mouse.

### Co-immunoprecipitation and Western Blot

Cells and dorsal root ganglia (DRGs) were harvested and lysed with ice-cold 0.5% NP-40 buffer containing protease inhibitors. The protein concentration of the lysates was determined using a BCA Protein Assay Kit (Beyotime). Immunoprecipitation was performed by incubating the lysates with rabbit anti-SARM1 (CST) antibody at 4°C for 4-6 hours, followed by incubation with Protein A/G PLUS-Agarose (Santa Cruz, sc-2003) at 4°C for an additional 4 hours. If a tagged protein (TAG) was co-expressed, TAG-beads were used instead of antibodies and Protein A/G PLUS-Agarose, and the incubation was carried out at 4°C for 4 hours. Following the immunoprecipitation, the Protein A/G Agarose or TAG-beads were washed three times with ice-cold NP-40 lysis buffer. The samples were then resolved by 12% SDS-PAGE, and proteins were transferred onto nitrocellulose membranes. After transfer, the membranes were blocked with 5% BSA and incubated with primary antibodies, including anti-β-actin as an internal control. The membranes were then incubated with HRP-conjugated secondary antibodies. Signals were detected using an enhanced chemiluminescence (ECL) system and imaged with CLINX Science Instruments.

### Measurement of NAD^+^ Levels

NAD^+^ levels in DRGs were measured using the Enhanced NAD+/NADH Assay Kit with WST-8 (Beyotime, S0176S), following the manufacturer’s provided protocol.

### Behavioral tests

8-week-old male mice were habituated to the environment for at least 2 days before testing. All tests were performed blindly. Von Frey Test: Mice were confined in boxes on an elevated metal mesh floor and habituated for at least 2 hours. Von Frey fibers with logarithmically increasing stiffness (0.02–2.56 g, North Coast) were applied to the central surface of hindpaws to determine mechanical thresholds of paw withdrawal using Dixon’s up-down method. Hargreaves Test: The radiant heat apparatus was used to measure paw withdrawal latency. Radiant intensity was adjusted to set the basal paw withdrawal latency of wild-type mice to 8–12 s. Each trial was repeated 3 times with 5-minute intervals, and a cutoff time of 20 s was set to prevent tissue damage. Hot Plate Test: Thermal sensitivity was assessed using a hot plate set at 50, 52, or 56°C. Latency to nociceptive responses (licking, paw shaking, or jumping) were measured, with cutoff times set at 60, 40, or 20 s, respectively, to avoid tissue damage. Tail-Immersion Assay: Mice were immobilized, and the tail tip (one-third of the length) was immersed in water at 48 or 52°C. Latency to tail withdrawal was measured, with cutoff times of 20 s and 10 s, respectively, for each temperature. Motor Function Test: The rotarod system assessed motor function by starting at 4 rpm and accelerating to 40 rpm over 5 minutes. Each trial was repeated 3 times at 20-minute intervals, and latency to fall was recorded. Paper-sticky Test: Mice were placed in a plexiglass chamber on an elevated glass floor. Adhesive tape was applied to the central surface of the hindpaws, and the response behaviors (shaking, scratching, or biting) were recorded over 5 minutes.

### AlphaScreen Assay

The assay was performed according to the manufacturer’s instructions (PerkinElmer). His-tagged PKM1 (2 μL at 300 nM) and biotinylated SARM1 (2 μL at 10 nM) were prepared in AlphaScreen reaction buffer (20 mM HEPES, pH 7.5, 100 mM NaCl, 0.1% BSA) and added to 384-well low-volume ProxiPlate microplates (10 μL per well). The mixture was incubated at room temperature for 60 minutes. After incubation, 1 μL of peptide was added, followed by another 60-minute incubation. Next, 2.5 μL of nickel-chelate acceptor beads were added and incubated for 60 minutes. Finally, 2.5 μL of Streptavidin-conjugated donor beads were added, and the plates were incubated for another 60 minutes. The signal was measured using a BioTek Synergy Neo2 multimode microplate reader (Agilent). Acceptor and donor beads were from the AlphaScreen Histidine (Nickel Chelate) Detection Kit (PerkinElmer).

### Determination of ALT, AST and BUN levels

The acute toxicity evaluation of the peptide was performed using the acute toxic class method with modifications. Mice were divided into peptide-treated (5 and 10 mg/kg) and control groups (n = 9). A single dose of saline-dissolved peptide (0.2 mL) was administered intraperitoneally to the treatment groups, while the control group received 0.2 mL of saline. Mice were monitored for 24 hours for signs of acute toxicity. Toxicity was assessed by observing animal behavior, followed by liver and kidney function tests. Blood samples (0.2 mL) were collected at 0, 1, 2, 4, 12, and 24 hours post-treatment via retro-orbital plexus into EDTA-coated vials. The serum was separated by centrifugation at 3000 rpm for 15 minutes at 4 °C. ALT, AST, and BUN levels were measured using commercially available detection kits (Solarbio Life Sciences) according to the manufacturer’s protocols.

## Data analysis

All experiments were performed with at least three biological replicates. Data are presented as means ± SD. Statistical significance was determined using the unpaired Student’s t-test (*P < 0.05, **P < 0.01, ***P < 0.001, ****P < 0.0001). Data analysis was conducted using GraphPad Prism 10.1.1.

## Data availability

The structural models used in this study are available in the Protein Data Bank under the following accession codes: SARM1 (7KNQ, 9L2F), and PKM1 (3SRF). Source data are provided in this paper.

---

**Supplementary Figure 1.**
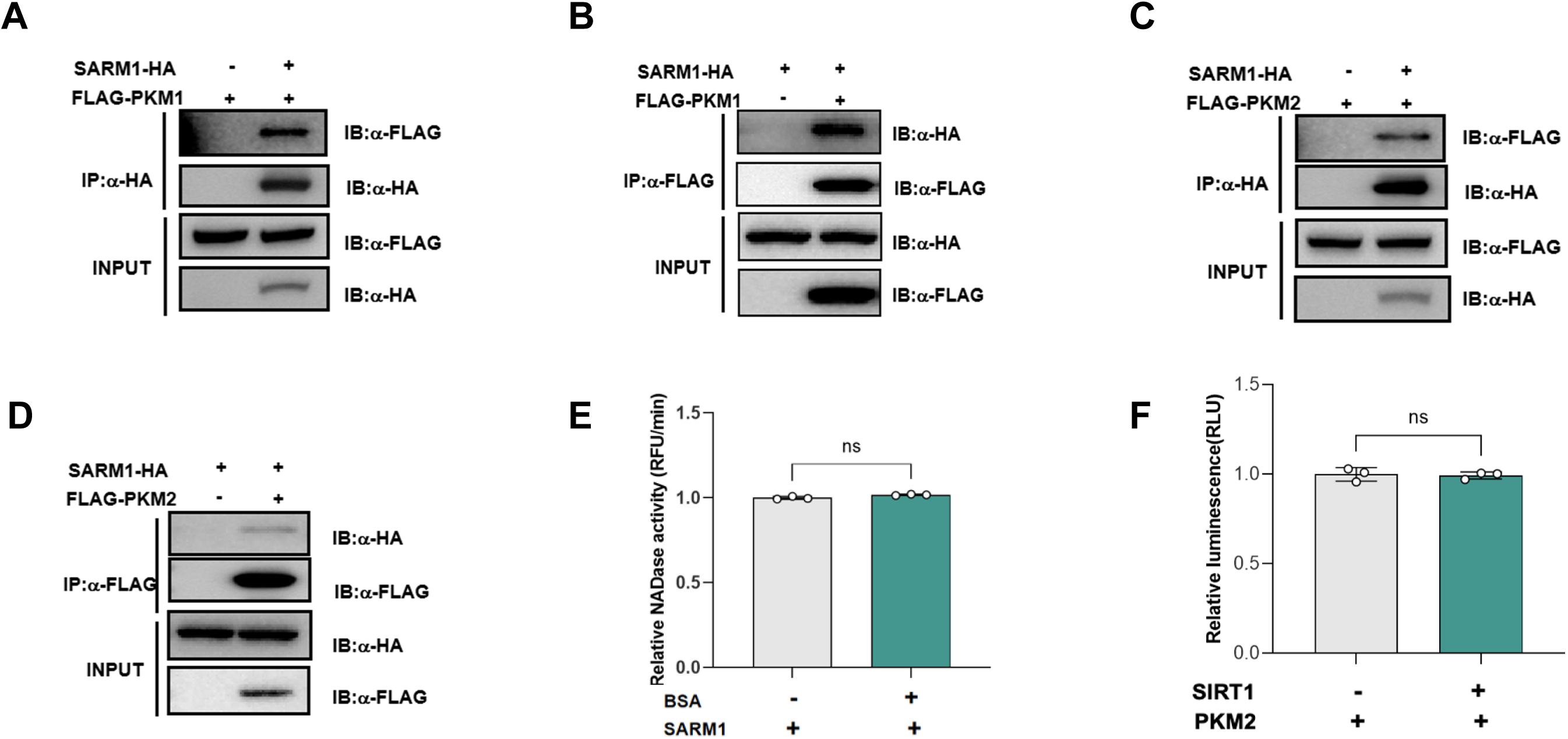
PKM directly binds SARM1 and allosterically enhances its NADase activity. (**A**) Interaction between overexpressed SARM1-HA and FLAG-PKM1 validated by HA immunoprecipitation in 293T cells. 293T cells were co-transfected with SARM1-HA (2 μg) and FLAG-PKM1 (2 μg) plasmids for 24 h. Cell lysates were incubated with HA-conjugated agarose beads. Immunoprecipitates were analyzed by Western blotting with anti-FLAG antibody. (**B**) Reverse IP of SARM1-HA and FLAG-PKM1 using anti-FLAG antibody followed by anti-HA blotting validates bidirectional binding. Input controls showed transfection efficiency. Data from two independent replicates with concordant results. (**C**) Interaction between overexpressed SARM1-HA and FLAG-PKM2 validated by HA immunoprecipitation in 293T cells directed as mentioned before. (**D**) Reverse IP of SARM1-HA and FLAG-PKM using anti-FLAG antibody followed by anti-HA blotting validates bidirectional binding. Input controls showed transfection efficiency. Data from two independent replicates with concordant results. (**E**) BSA control validates the specificity of the SARM1 NADase activity assay. Recombinant SARM1 (6 µg/mL) was incubated with NAD^+^ (2000 μM), NMN (500 µM) and PC6 (50 µM) in the presence or absence of bovine serum albumin (BSA, 1 mg/mL). BSA supplementation showed no significant effect on SARM1-dependent NAD^+^ degradation (ns: p > 0.05, unpaired t-test). Data confirm that SARM1 activity measurements are independent of nonspecific protein interactions. Error bars: means ± SD (n = 3). (**F**) SIRT1 does not modulate PKM2 pyruvate kinase activity. Recombinant PKM2 (100 nM) was incubated with phosphoenolpyruvate (2 mM) and ADP (1 mM) with or without SIRT1 (200 nM). SIRT1 exhibited no regulatory effect on PKM2 catalytic activity (ns: p > 0.05). Results exclude potential crosstalk between NAD^+^-dependent deacetylases and PKM2 function. Data from two independent experiments. Error bars: means ± SD (n = 3).

**Supplementary Figure 2.**
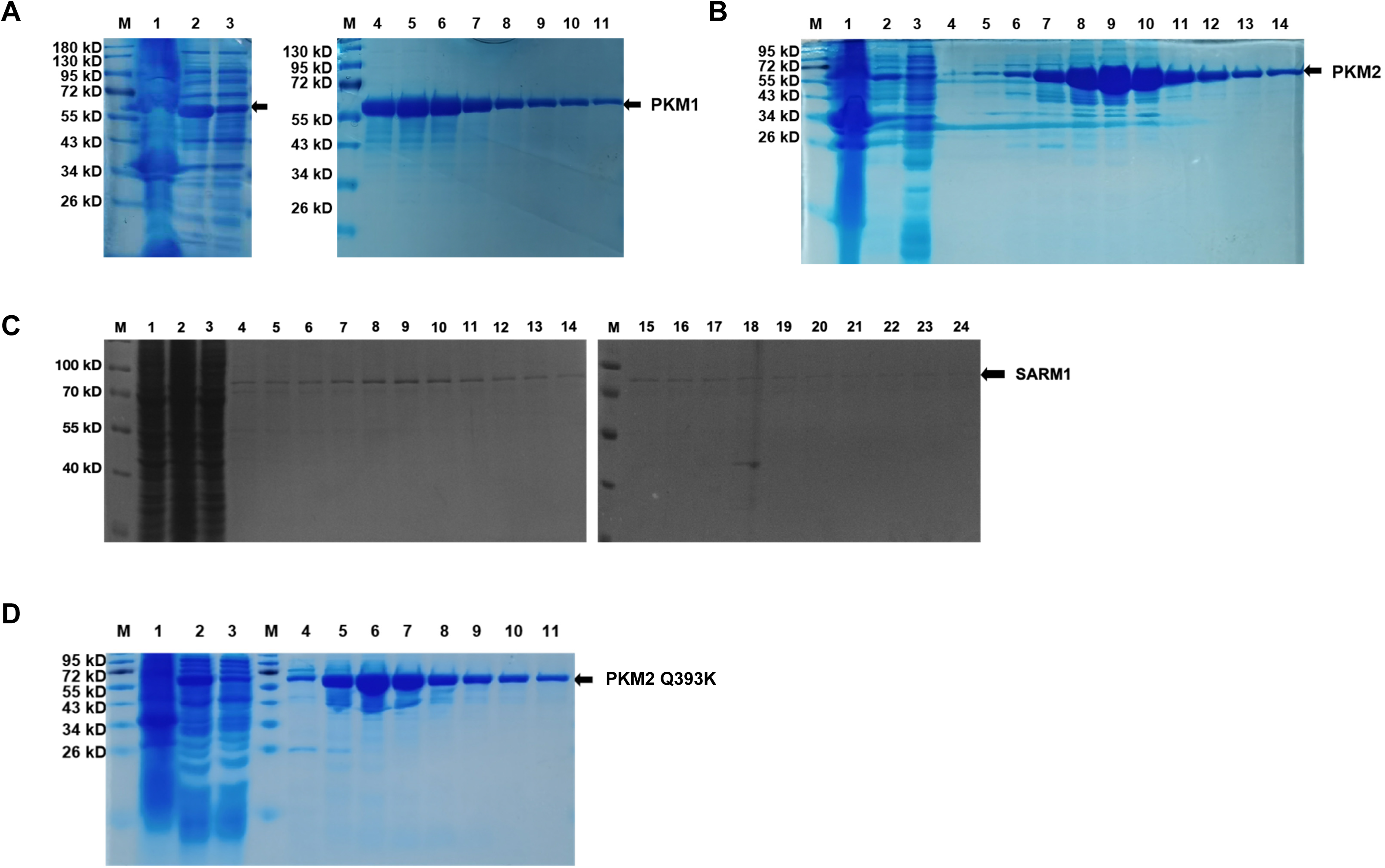
Expression and purification of PKM1, PKM2, SARM1 and PKM2 Q393K. PKM1(**A**), PKM2 (**B**), SARM1 (**C**), PKM2-Q393K (**D**) protein preparations. Presented are SD-PAGE analyses of the metal-chelate chromatography. Arrows indicate PKM1 (58 kDa), PKM2 (58 kDa), SARM1 (72 kDa) and PKM2-Q393K (58 kDa).

**Supplementary Figure 3.**
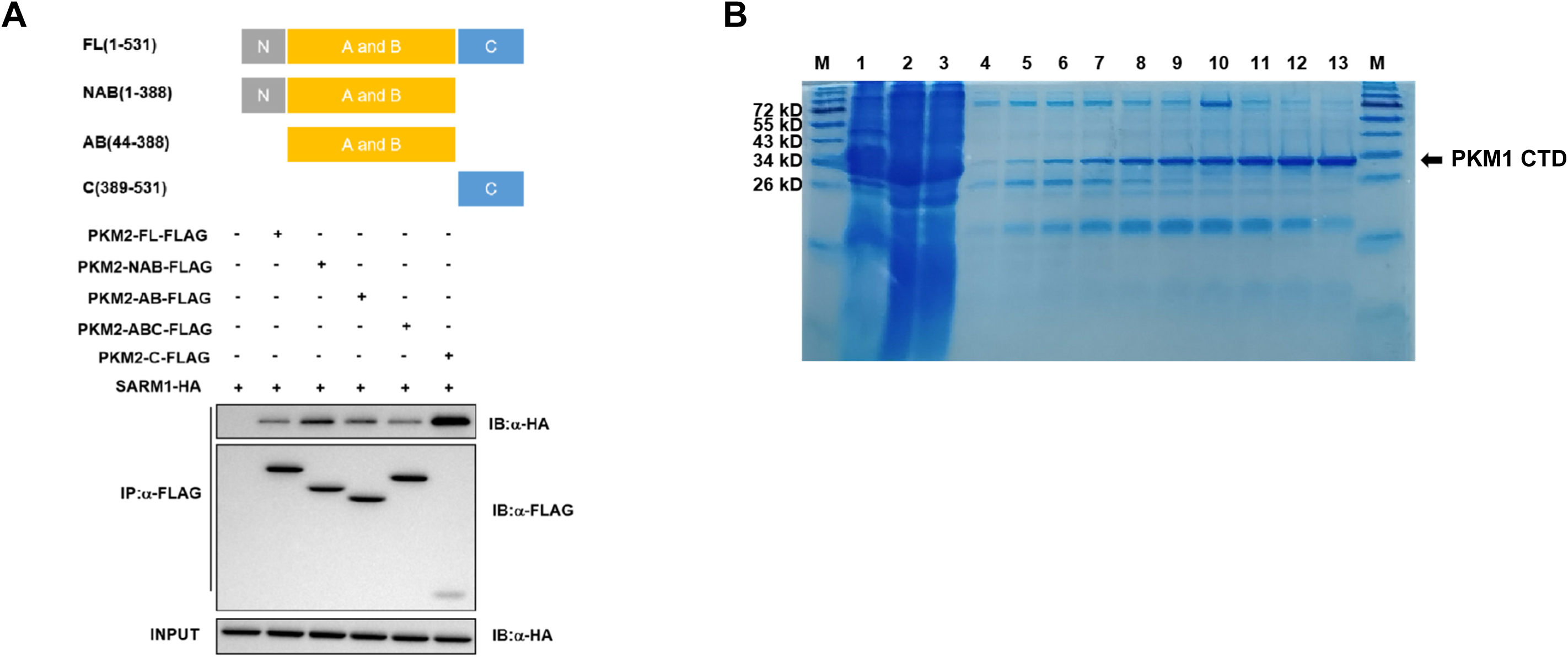
PKM C-terminal domain mediates direct binding to the SARM1 SAM-TIR region. (**A**) The C-terminal of PKM2 strongly interacts with SARM1 and enhances its enzymatic activities. 293T cells were co-transfected with full-length SARM1-HA (2 μg) and PKM2-FLAG truncation mutants (2 µg). FL: full-length PKM2 (residues 1-531); NAB: the N-terminal domain, A and B domain of PKM2 (residues 1-388); AB: the A and B domain of PKM2 (residues 44-388); C: the C-terminal domain of PKM2 (residues 389-531). Despite the lower precipitation efficiency of PKM2 C-terminal protein, SARM1 showed the strongest enrichment when co-precipitated with the C-terminal truncation. Input lanes (10% total lysate) showed protein expression levels. (**B**) PKM1 C-terminal domain protein preparations. Presented are SD-PAGE analyses of the metal-chelate chromatography. Arrow indicates PKM1-CTD (34 kDa).

**Supplementary Figure 4.**
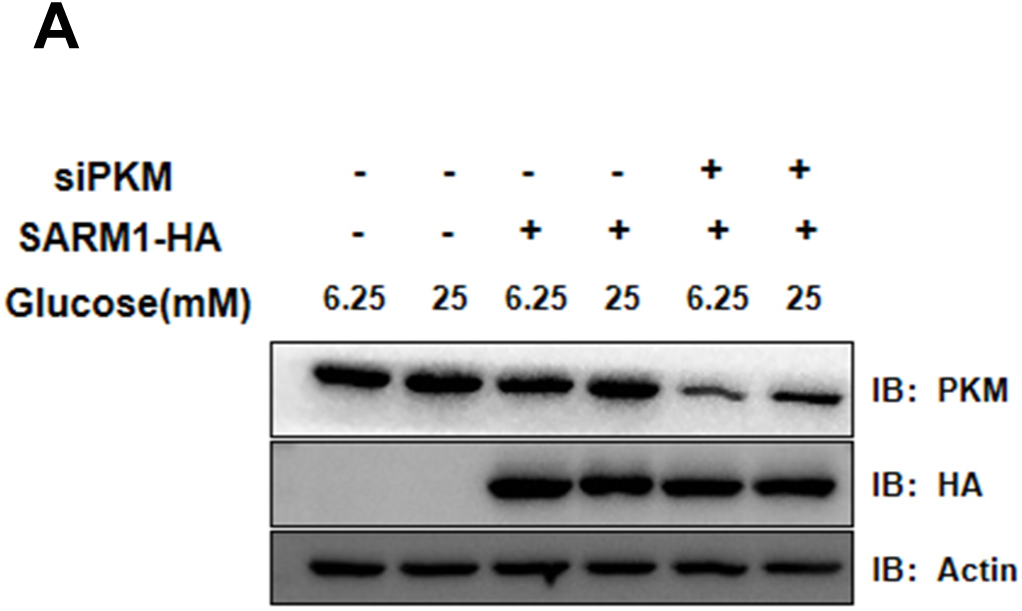
In vivo verification of PKM knockdown efficacy. (**A**) siRNA-mediated knockdown of PKM in 293T cells. Immunoblot analysis showing the reduction of PKM protein levels following siRNA transfection. Actin serves as a loading control.

**Supplementary Figure 5.**
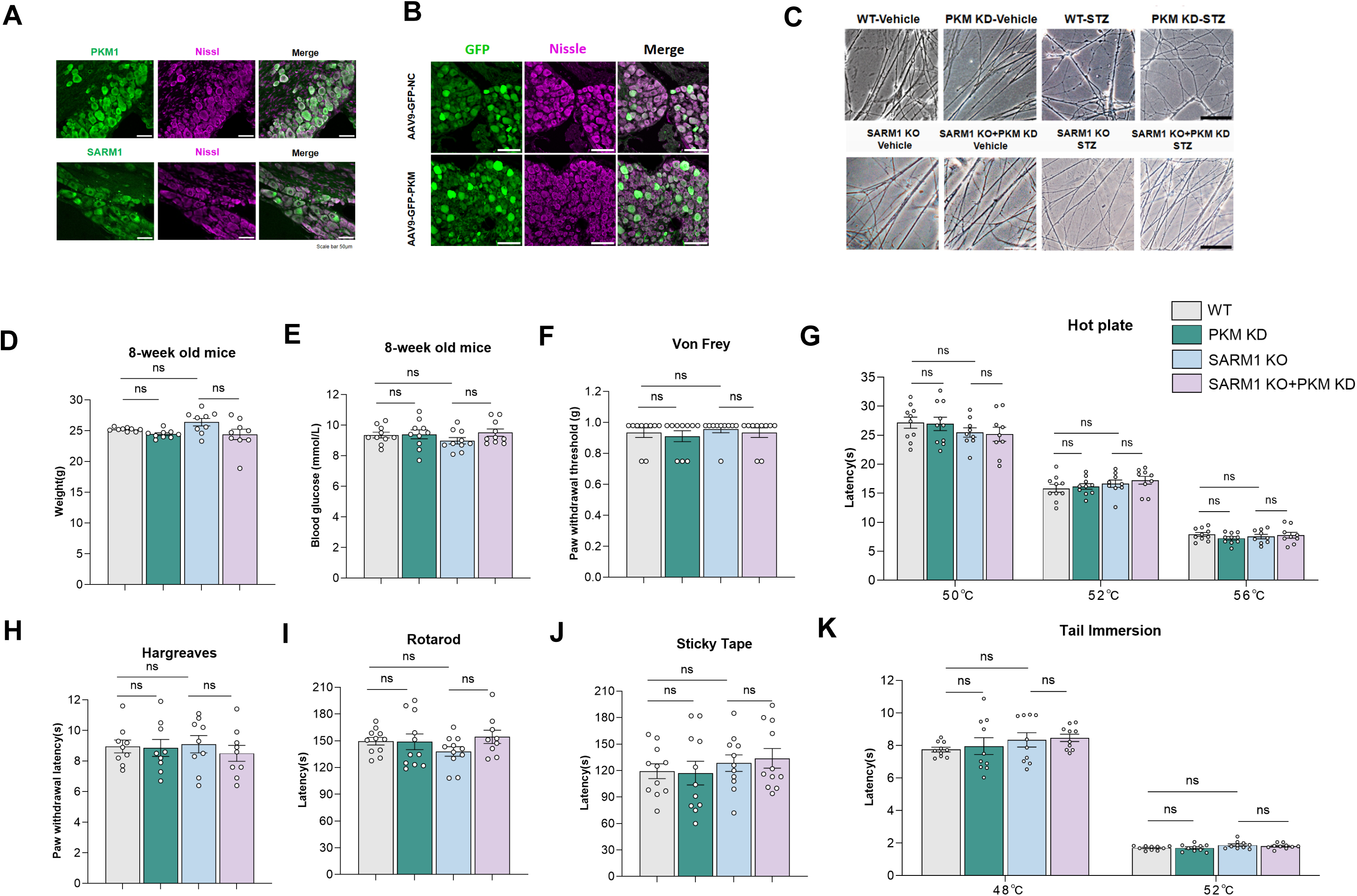
SARM1 KO and PKM KD mice exhibited normal basal motor and sensory function. **(A)** Co-localization of PKM1 and SARM1 with Nissl in DRGs. Scale bar 50 μm. (**B**) GFP fluorescence detection in DRGs frozen sections to confirm PKM KD efficiency. (**C**) Representative phase-contrast images showing DRG neuronal axonal degeneration caused by the STZ-induced diabetic neuropathy in WT, PKM KD, SARM1 KO and SARM1 KO and PKM KD mice. Quantification of the axonal degeneration (right). n = 5 mice per group; scale bar: 50 μm. (**D, E**) WT, PKM KD, SARM1 KO, and SARM1 KO and PKM KD mice exhibit similar body weight and blood glucose levels at 8 weeks of age. (**F**) Mechanical pain sensation was measured through the von Frey test. (**G**, **H** and **K**) Thermal sensation was assessed through hot plate assay (**G**), Hargreaves assay (**H**), and tail immersion assay (**K**). (**I**) Motor function was tested through Rotarod assay. (**J**) The light-touch sensation was tested through the paper-stick test. n = 9 to 12 mice per group. (means±SD; Student’s t-test, *p<0.05, **p<0.01, ****p<0.0001)

**Supplementary Figure 6.**
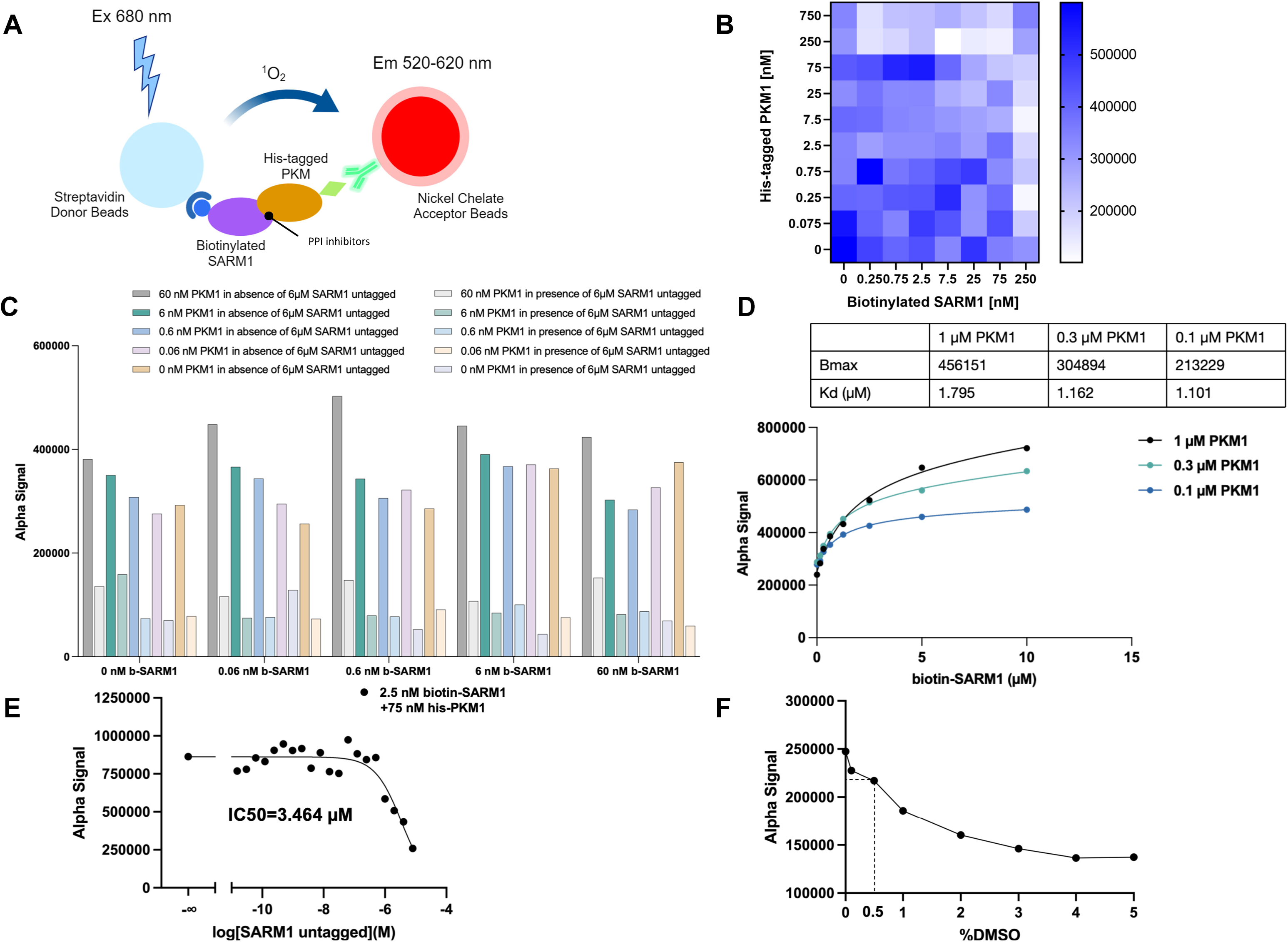
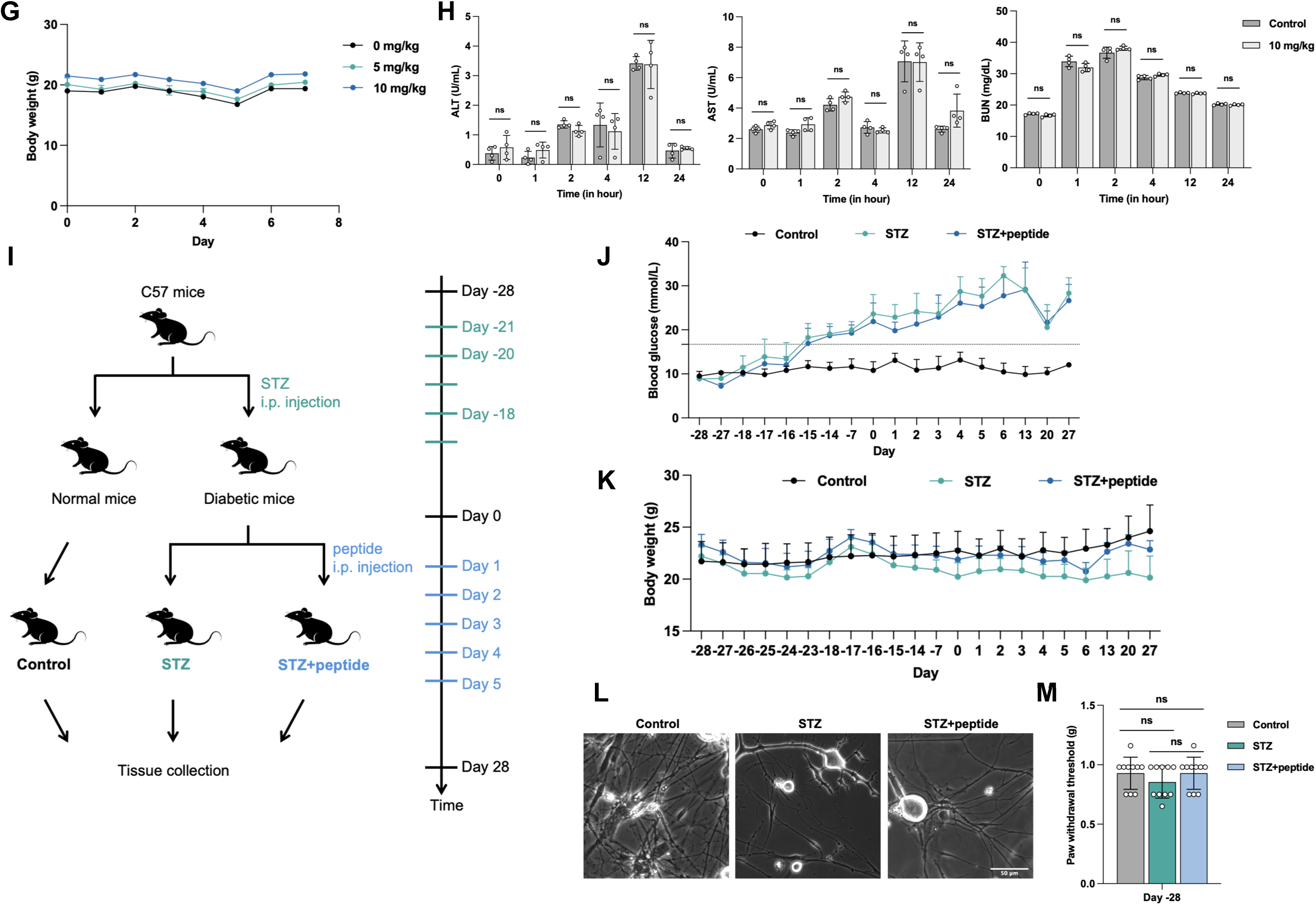
AlphaScreen platform optimization, acute toxicity profiling, and efficacy evaluation in STZ-induced diabetic mice. (**A**) Schematic representation of the AlphaScreen assay principle. (**B**) Cross-titration analysis of biotinylated SARM1 binding to His-tagged PKM1. Based on the signal-to-noise ratio, 2.5 nM biotin-SARM1 and 75 nM His-PKM1 were selected for subsequent experiments. (**C**) Comparison of signal derived from the biotin-SARM1:his-PKM1 PPI assay. (**D**) Saturation binding curves for determining the dissociation constant (Kd) of the SARM1-PKM1 interaction. The Kd values at different PKM1 concentrations (0.1, 0.3, 1 µM) are shown. (**E**) A competitive binding assay using untagged SARM1 to confirm the specificity of the SARM1-PKM1 interaction (IC50 = 3.464 µM). (**F**) Effect of DMSO concentration on the AlphaScreen assay signal. A final concentration of 0.5% DMSO was chosen for all subsequent experiments. (**G**) Effect of peptide administration at doses of 0, 5, and 10 mg/kg body weight on body weight changes in mice over time. (**H**) Assessment of liver (ALT, AST) and kidney (BUN) function biomarkers in mice treated with 10 mg/kg peptide. Serum dilution ratios had no significant impact on biomarker measurements. Data are presented as mean ± SD (n = 9). (**I**) Experimental timeline for STZ-induced diabetic mouse model and peptide intervention. C57 mice were injected intraperitoneally with STZ from day -21 to day -17 to induce diabetes. From day 1 to day 5, diabetic mice received daily intraperitoneal injections of the peptide. The effect of peptide treatment was evaluated up to day 27. (**J**) Random blood glucose levels were monitored throughout the study (-28 to 28 days). Mice developed hyperglycemia (≥16.8 mmol/L) starting from day -15, confirming the establishment of the diabetic model. Control: normal mice; STZ: STZ-induced diabetic mice; STZ+peptide: peptide-injected STZ-induced diabetic mice. (**K**) Body weight monitoring throughout the study (-28 to 28 days). STZ-treated mice showed weight loss compared to controls. (**L**) Axonal integrity in DRG neurons. Control: normal mice; STZ: STZ-induced diabetic mice; STZ+peptide: peptide-injected STZ-induced diabetic mice. Representative phase-contrast images are shown. n = 3 mice per group; scale bar: 50 μm. (**M**) Baseline von Frey test results at day -28 for the control, STZ, and STZ+peptide groups. All groups had similar mechanical sensitivity thresholds (∼0.98 g), with no significant differences observed (ns: p > 0.05). Error bars: means ± SD (n = 10).

## Notes

### Competing Interest Statement

The authors have declared no competing interest.

